# Short translational ramp determines efficiency of protein synthesis

**DOI:** 10.1101/571059

**Authors:** Manasvi Verma, Junhong Choi, Kyle A. Cottrell, Zeno Lavagnino, Erica N. Thomas, Slavica Pavlovic-Djuranovic, Pawel Szczesny, David W. Piston, Hani Zaher, Joseph D. Puglisi, Sergej Djuranovic

**Affiliations:** Department of Cell Biology and Physiology, Washington University School of Medicine, 600 South Euclid Avenue, Campus Box 8228, St. Louis, MO 63110, USA; Department of Structural Biology, Stanford University School of Medicine, Stanford, California 94305– 5126, USA; Department of Applied Physics, Stanford University, Stanford, California 94305– 5126, USA; Department of Biology, Washington University, St Louis, Missouri 63105, USA; Institute of Biochemistry and Biophysics Polish Academy of Sciences, Department of Bioinformatics, Warsaw, Poland

## Abstract

It is generally assumed that translation efficiency is governed by translation initiation. However, the efficiency of protein synthesis is regulated by multiple factors including tRNA abundance, codon composition, mRNA motifs and amino-acid sequence^1–4^. These factors influence the rate of protein synthesis beyond the initiation phase of translation, typically by modulating the rate of peptide-bond formation and to a lesser extent that of translocation. The slowdown in translation during the early elongation phase, known as the 5’ translational ramp, likely contributes to the efficiency of protein synthesis ^5–9^. Multiple mechanisms, which could explain the molecular basis for this translational ramp, have been proposed that include tRNA abundance bias^6,9^, the rate of translation initiation^10–15^, mRNA and ribosome structure ^11,12,14,16–18^, or retention of initiation factors during early elongation events ^19^. Here, we show that the amount of synthesized protein (translation efficiency) depends on a short translational ramp that comprises the first 5 codons in mRNA. Using a library of more than 250,000 reporter sequences combined with *in vitro* and *in vivo* protein expression assays, we show that differences in the short ramp can lead to 3 to 4 orders of magnitude changes in protein abundance. The observed difference is not dependent on tRNA abundance, efficiency of translation initiation, or overall mRNA structure. Instead, we show that translation is regulated by amino-acid-sequence composition and local mRNA sequence. Single-molecule measurements of translation kinetics indicate substantial pausing of ribosome and abortion of protein synthesis on the 4^th^ or 5^th^ codon for distinct amino acid or nucleotide compositions. Introduction of preferred sequence motifs, only at the exact positions within the mRNA, improves protein synthesis for recombinant proteins, indicating an evolutionarily conserved mechanism for controlling translational efficiency.

## Main

The efficiency of protein synthesis is governed by the rates of translation initiation, elongation and to a lesser extent termination ^1–4^. Studies have investigated potential factors that contribute to the protein synthesis efficiency using both endogenous genes and reporter sequences by focusing on tRNA abundance, amino acid sequence or both mRNA sequence and structure ^6,9,12,17,19–27^. Several conflicting models have been proposed regarding the role of codon distribution at the N-terminus as well as the local mRNA structure around the translation start sites on the efficiency of protein synthesis ^6,9,11,12,25^. Reduced abundance of tRNAs coding for N-terminal residues of proteins may play a crucial role in slowing down initial stages of translation elongation, which could in turn decrease the cost of gene expression by reducing the probability of ribosome jamming during translation ^6,9,15^. Such a translational ramp would be beneficial in preventing detrimental collision-dependent abortion of protein synthesis ^9,27,28^. Some of these effects can be rationalized by the presence of mRNA structure structural elements within the first 5 to 16 codons ^10–12,29,30^. In addition, interactions between the nascent peptide and the exit tunnel of the ribosome appear to play an important role in dictating the rate of peptidyl transfer during these early elongation events ^16–18,31^. However a unifying mechanism that could explain the interplay between protein-synthesis yield and rates of early elongation events remains unknown. Here, we present data that strongly suggest that the mRNA and the encoded protein sequences of the first five codons are key in dictating the efficiency of protein synthesis.

To decipher how mRNA sequence and the encoded nascent peptide influence the efficiency of protein synthesis, we focused on the region surrounding +10 nucleotide position of a GFP-reporter sequence. Previously, this region has been implicated in regulating protein expression levels ^11,12,25^. We created a library of an otherwise fully-optimized eGFP gene with an insertion of 9 random nucleotides after the second codon (Fig 1A). We were able to obtain a total of 259,134 unique sequences out of the 262,144 possible synthetic eGFP constructs. These were nearly identical save for the 3^rd^, 4^th^ and 5^th^ codons of the open reading frame. These three codons coded for 9261 different tri-peptide sequences including truncated peptides due to the presence of one or more stop codons in the variable region. The library was cloned into the low-copy pBAD plasmid vector and expressed using arabinose induction in *E. coli* DH5-alpha cells. To score global expression of the eGFP variants, bacterial cells were FACS sorted into five bins (Fig 1B) with a difference in relative fluorescence units (RFUs) of almost four orders of magnitude between bin one and bin five (Fig 1C, fluorescence difference >600-fold). Interestingly, this difference is higher than previously reported for expression of 14,000 variants of super-folder green fluorescent protein (sfGFP) with randomized promoters, ribosome binding sites, and first 11 codons ^11^, or when 94 % of the eGFP protein was recoded using synonymous codons ^12^. These results indicated that the overall expression of the protein could be significantly changed as a result of differences in the N-terminal sequence that encompasses the first 5 amino acids of the protein, in other words nucleotides 7-15 of the open-reading frame (ORF) corresponding to the first ribosome footprint after initiation (Supplementary Fig 1).

**Figure 1.**
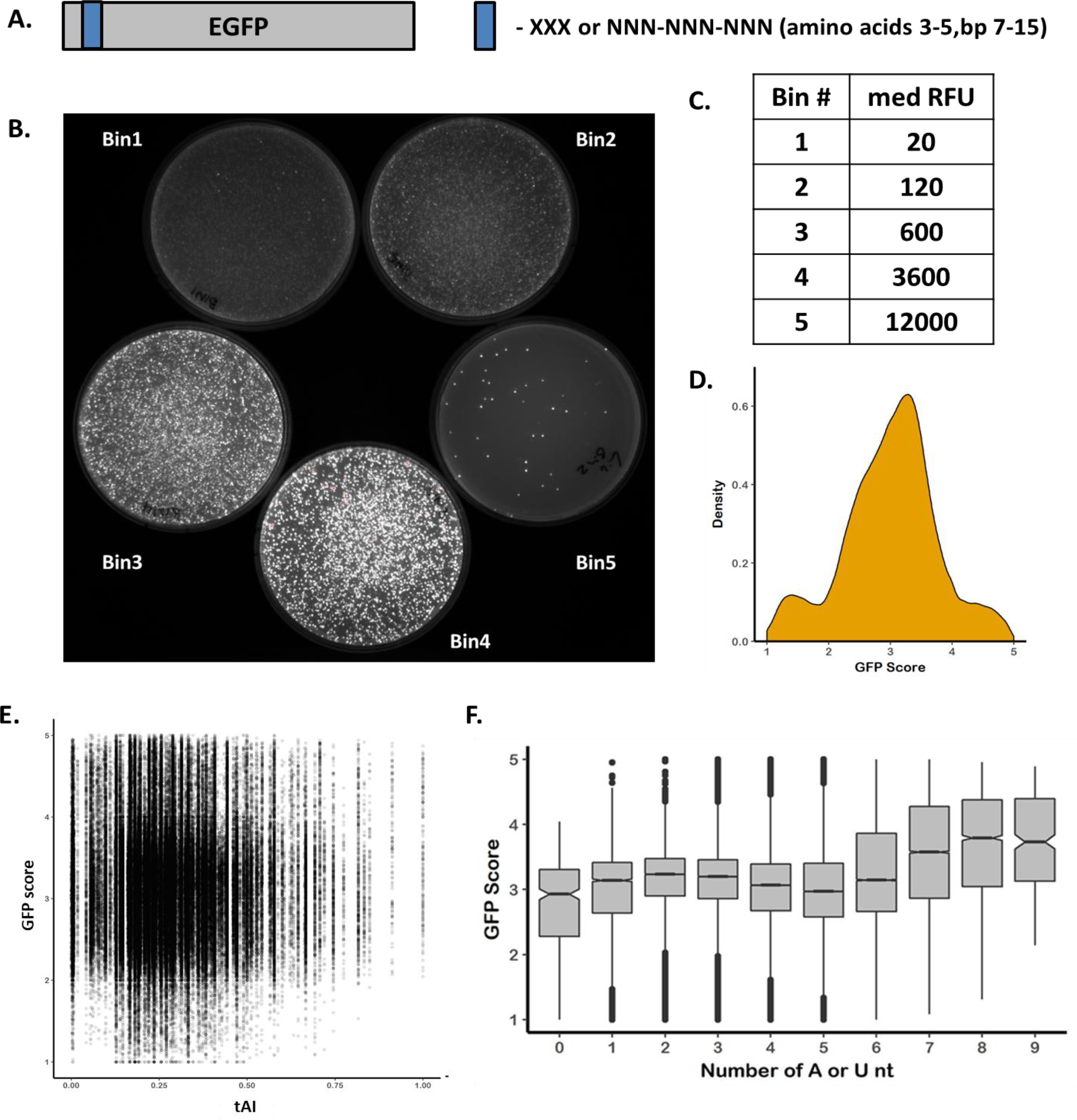
A. Scheme of the reporter system to test influence of the first 5 amino acids and mRNA sequence of the first ribosome footprint. Sequences of 9 random nucleotides were introduced into eGFP reporter coding for all amino acid and stop codon possibilities in the positions from 7-15 nucleotide coding for amino acids 3, 4 and 5 in protein. Library was introduced in a single (low) copy pBAD plasmid with arabinose inducible promoter. B. Image of arabinose induced E. coli colonies separated using fluorescence-activated cell sorting (FACS) into 5 bins. Each bin represents approximately 24% of the whole cell population depending on EGFP expression with exception of bin 5. Bin 5 represents 2.5% of the E.coli cells with highest EGFP expression based on relative fluorescence values (RFUs). C. Table of relative average fluorescence values for colonies in five separated bins. Wild type eGFP expression is approximately 250 RFUs. D. Distribution of the plasmid reads based on the GFP score. GFP score represents distribution value for each independent sequence in 5 bins. E. No correlation is observed between tRNA abundance (tAI –tRNA abundance index) and the expression of eGFP variants based on GFP scores. F. Influence of local mRNA structure on expression of eGFP 9nt library. GFP score distribution value is plotted in correlation with the number of A or U nucleotides in 9nt randomized sequence. Boxplot whiskers indicate the furthest datum that is 1.5*Q1 (upper) or 1.5*Q3 (lower).

To assign expression level for each sequence variant, plasmids were isolated from each bin, normalized by DNA amounts and sequenced (Supplementary Fig 2). Resulting counts for each variant were normalized and the GFP score was calculated to represent the weighted distribution value for each independent sequence over five FACS sorted bins (Fig 1D and Supplementary Fig 2). A GFP score close to 1 indicates sequences with low eGFP expression for which the majority of sequencing counts associated with bin 1; a GFP score of 5 specifies sequences that are highly expressed and coupled mostly with bin 5 (Supplementary Tables 1-5). On average, the complete library had a score slightly over 3 with most of the sequences distributed between bins 2, 3 and 4. GFP scores were consistent and reproducible across the 213708 constructs in the library with at least 10 reads, with a Pearson correlation of 0.8 among biological replicates (Supplementary Fig 2). Since *amber* stop codon (UAG) suppression in DH5α is highly efficient (75-95%)^32^ we used this feature of the *E.coli* DH5α cells to compare eGFP variants with amber stop codon in randomized positions 3, 4 or 5 with other stop codons (opal-UGA, ochre-UAA, Supplementary Fig. 3). While variants with *ochre* and *opal* stop codons distributed between GFP scores of 1 and 2, distribution of the constructs with an amber stop codon followed the distribution of the library without encoded stop codons (188,703 variants). As such amber suppressor tRNA (supE44)^32^ that codes for tRNA^Gln^_CUA_ and leads to Gln incorporation at UAG codon served as an additional control for the codon-anticodon interaction and efficiency of protein synthesis (Supplementary Fig 3).

To test whether eGFP reporter levels depend on tRNA abundance or rare codons at the start of the coding sequence, we compared the distribution of the GFP scores of all library variants to these features (Fig 1E and Supplementary Fig 3). We did not find any obvious correlation of GFP scores with tRNA abundance (Fig 1E), measured by tRNA adaptation index (tAI)^33^, or when rare Arg, Ile or Leu codons were at the 3^rd^, 4^th^ and/or 5^th^ codon (Supplementary Fig 3). We also found no correlation with the amino-acid chemical properties such as overall charge or hydrophobicity of the encoded tri-peptides (Supplementary Fig 4). There was also no correlation between GFP score and plasmid abundance in the unsorted cells (Supplementary Fig 2). We found that GFP score correlated moderately with the AT content of variable region similar to the so-called downstream box element (Fig 1F and Supplementary Fig 5)^10,14,34,35^. Indeed, eGFP variants that harbored 6-9 A or T nucleotides at positions +7 to +15 had on average better expression than the rest of the library variants. This was further confirmed with more thorough analysis of the library sequences divided into four buckets defined by GFP score (Supplementary Fig 5). Each bucket represented X≤GFP score<(X+1), where X was 1-4. Sequences motif analysis of variants with highest GFP scores (GFP score>4) indicated slight AT bias; however, there was not strong bias against GC rich sequences (Supplementary Fig 5). Sequences that were moderately expressed had a more or less random distribution of GC nucleotides, with a slight increase in C nucleotides for low expressed sequences (1≤GFP score<2). This could potentially be explained by the C-rich codons for proline and previously described proline stalling during translation ^36–38^. Taken together, these analyses indicate that local mRNA sequence and potentially base-pairing stability of nucleotides +7 to +15 influence expression of the protein.

Given that AT-richness correlated slightly with eGFP variants with higher expression, we asked whether certain sequences influenced expression of eGFP. Simple analyses of eGFP variants with expression scores over 4 indicated slight preference for certain amino acids in positions 3, 4 or 5 (Supplementary Fig 6). Using a motif-scanning approach, we identified motifs that were enriched in eGFP variants with a score greater than 4, when compared to poorly expressed variants (GFP score <3). Among several hexanucleotide motifs that were identified, the two most significantly enriched RNA motifs (enrichment ratio of >9 and p value < 1E-5) were AADTAT (D stands for not C, Figure 2A) and AAVATT (V stand for not T, Figure 2B). During decoding, these motifs code for lysine (K) or asparagine (N) and tyrosine (Y) or isoleucine (I), as first and second amino acids, respectively. Intriguingly, all eGFP variants with combination of K|N-Y|I amino acids regardless of their synonymous codons had on average a GFP score of 4.2±0.4, arguing for possible amino acid contribution for higher expression (Supplementary Fig 7). These same amino acids were identified as occurring more frequently in eGFP variants with high score (>4) compared to those with low score (Supplementary Fig 6). Analyses of the positional bias of hexanucleotide motifs (Fig 2C) revealed that K|N-Y|I amino acid combination on average had higher GFP scores than any other amino acid combination encoded by the same hexanucleotide motifs in a randomized 9 nucleotide sequence (Fig 2D-2G). We observed also preference for certain amino acids at position 3 when K|N-Y or K|N-I motifs were in position 4|5 (Supplementary Fig 8). Analysis of the influence of K, N, I and Y isoacceptor tRNAS for indicated rather small differences between tested codons. Tendency for reduction in GFP scores was observed for codons with G or C nucleotides and when motifs were shifted to position 4 (Fig 2E and 2I, Supplementary Fig 7). These analyses suggest that both amino acid and nucleotide composition at the N-terminus and beginning of the coding sequence, respectively, contribute to the overall efficiency of protein synthesis.

**Figure 2.**
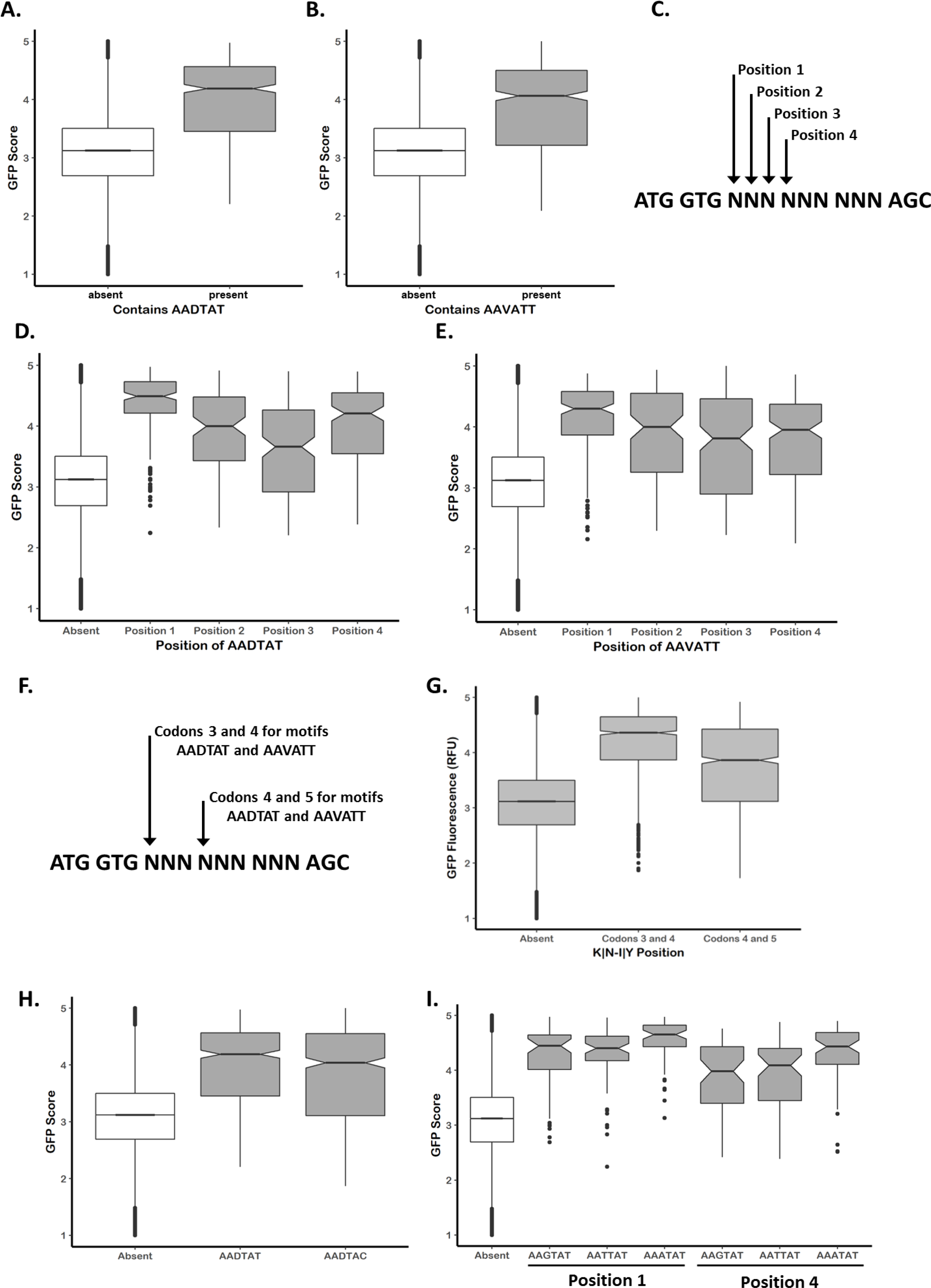
A. and B. enrichment analyses of sequenced constructs with average GFP Score of ≥4.0 results in two motifs with DNA sequence AADTAT and AAVATT, or amino acid sequence K|N-Y and K|N-I, respectively. Average GFP score of all sequences with two motifs (present) is compared to the rest of library (absent). C. Scheme of analysis of the GFP scores for two motifs by movement one nucleotide at the time. Position 1 and position 4 code for K|N-Y and K|N-I amino acid motifs as codons 3 and 4 or 4 and 5, respectively. D. and E., analysis of average GFP scores for two sequences motifs based on their position in 9nt randomized sequence indicates potential amino acid dependence. Average GFP score is compared to the rest of library (absent). F. Scheme of analysis of overall influence of amino acid sequence when motifs code for amino acids in positions 3 and 4 or 4 and 5, respectively. G. Analysis of overall influence of amino acid sequence of motif K|N-I|Y in positions 3, 4 and 5. Average GFP score for motifs is compared to the rest of library (absent). H. and I. Analysis of the influence of degenerate codons for Tyr or Asn and Lys on the GFP score of AADTAT motif, respectively. All analyzed sequences with stop codons were filtered out to represent average coding library (absent). Comparison is shown vs all the coding constructs in the library. Boxplot whiskers indicate the furthest datum that is 1.5*Q1 (upper) or 1.5*Q3 (lower).

We next probed the effects of mRNA and protein stability on the expression of the reporter protein *in vitro* and *in vivo*. In particular, we compared the expression of wild type eGFP (WT) and AADTAT hexanucleotide variants (Mp1-Mp4, Supplementary Fig 9) by western analysis (Fig 3A, Supplementary Fig 9), kinetics of *in vitro* protein synthesis (Supplementary Fig 10) and endpoint eGFP fluorescence for both *in vivo* and *in vitro* experiments (Fig 3B). Results from both *in vivo* and *in vitro* experiments confirmed our FACS and bioinformatics analyses of the library, for which Mp1-Mp4 eGFP variants displayed higher expression levels than WT construct. We noted that *in vitro* expression of Mp1-Mp4 constructs showed moderately higher levels (range 3-10 fold higher than WT) when compared to the *in vivo* expression in *E. coli* BL21 cells (3-6 fold higher than WT), suggesting some contribution of protein degradation and mRNA stability to the observed difference in protein yields. However, these results also indicated that protein and mRNA stability do not strongly contribute to alterations in eGFP expression driven by amino acid identities in position 3-5 and ORF nucleotides 7-15 present in the Mp1-Mp4 constructs. In addition, expression of WT eGFP and two randomly picked reporter constructs coding for NCT (GFP score of 3.04±0.40, 5 fold higher expression than WT) and LQI (GFP score of 2.67±0.40, 3 fold higher expression than WT) in positions 3-5 maintained the difference in expression ratio regardless of the change in the 2^nd^ amino acid (Fig 3C) or when a different *E. coli* strain was used for expression (Supplementary Fig 11). Finally, changing the starting codon (AUG) to near-cognate start codons (GUG, UUG) in three different eGFP variants resulted in overall reduction of eGFP expression as observed previously ^39^, but the relative expression difference between the three sequences was unaffected by the start codon (Supplementary Fig 12). As such, we deduced that expression differences of analyzed eGFP variants were not driven by overall protein or mRNA stability (*in vitro* or *in vivo*) or character of the 2^nd^ amino acid (N-end rule)^40^. The difference in the ratio of expression for tested eGFP reporters was maintained despite usage of different *E. coli* strains (K - DH5α, W3110, XAC or B – BL21(DE3)) or reduced efficiency of start codon recognition during initiation on near-cognate start sites. These results suggest that observed differences in expression of reporter variants are driven by composition and position of nucleotides and amino acid sequence between 7-15 nucleotide and 3-5 amino acid, respectively.

**Figure 3.**
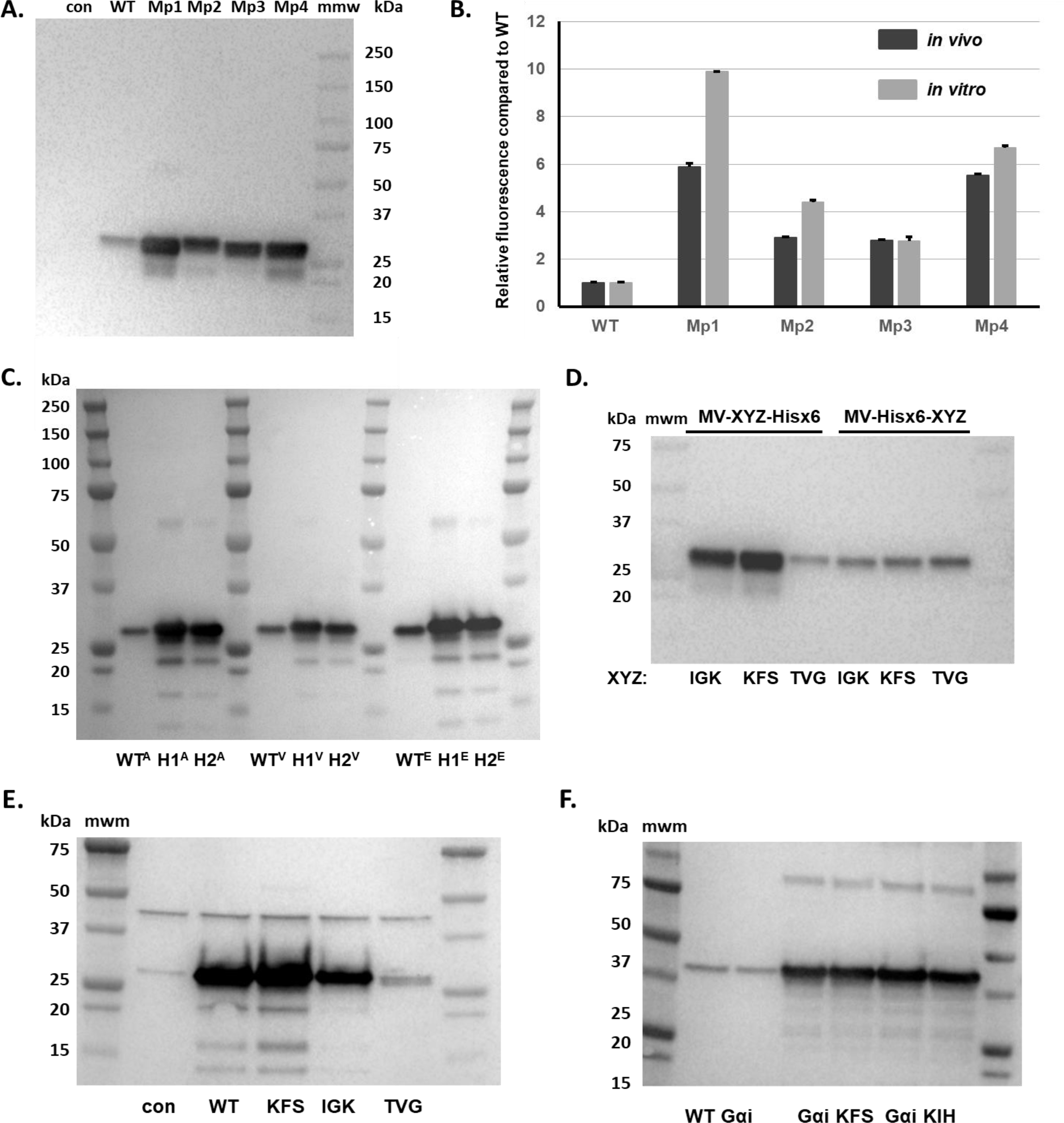
A. Western blot analysis of NEB Pure Express in vitro expression of eGFP constructs with motif AAD TAT in different positions coding for amino acids 3, 4 and 5. M1p1 indicates motif1 in position 1, M1p2 indicates motif1 in position 2, M1p3 indicates motif1 in position 3 and M1p4 indicates motif1 in position 4, where insertion positions are defined as in Figure 2C. Wild type eGFP (WT) control is indicated. 5% of *in vitro* translation reaction is analyzed. B. Relative eGFP fluorescence from in vitro and in vivo expression of eGFP constructs with AADTAT motif in different position compared to the wild type eGFP construct. NEB Pure Express expression system and pBAD single copy vector in BL21 cells were used for in vitro and in vivo expression, respectively. C. Western blot analysis indicates that N-terminal rule does not influence expression of eGFP variants from pBAD single copy vector in E. coli Top10 cells. Two high expression variants H1 (NCT) and H2 (LQI) and WT eGFP constructs are indicated. Letter in superscript indicates amino acid in the second position (A-alanine, V-valine, E-glutamic acid). D. Positional bias in controlling expression of eGFP constructs with different amino acids in position 3, 4 and 5. Western blot analysis of NEB Pure Express in vitro expression of eGFP constructs with sequence XYZ (KFS, IGK and TVG, respectively) as amino acids 3(X), 4(Y) and 5(Z) followed by 6xHis tag (MV-XYZ-6xHis) or as amino acids 9(X), 10(Y) and 11(Z) preceded by 6xHis tag (MV-6xHis-XYZ). eGFP antibody (J8, Promega) is used to visualize expression of eGFP. 5% of in vitro translation reaction is analyzed. E. and F. simple insertion of different amino acids in recombinant mEOS2 or human Gαi protein constructs can modulate their expression in E.coli BL21 cells, respectively. Wild type (WT) and control samples as well as amino acids in position 3, 4 and 5 for variants of mEOS2 and Gαi proteins are indicated. mEOS2 and Gαi constructs were cloned in pet16b and pBAD vector as C-terminally His-tagged proteins. Proteins were visualized based on their C-terminal 6-Hist tag using Penta-His (Qiagen) antibody. GFP antibody (J8, Promega) is used to visualize expression of eGFP and Biorad Precision Plus marker is indicated in all images. Same amount of the E.coli cells (OD600) was used for Western Blot analysis of in vivo expression of different reporter constructs.

To test whether our results depend on the position of an amino acid or nucleotide motif with respect to the start codon, and are therefore independent of the rest of the reporter sequence, we created several new reporters. First, we picked three eGFP variants with somewhat different expression levels (KFS for high, GFP Score of 4.90±0.02; IGK for medium, GFP Score of 3.01±0.44; and TVG for low expression, GFP score 1.93±0.21) and inserted 6xhistidine tag between the variable sequence and eGFP sequence (creating constructs MV-XYZ-6xHis-eGFP, where XYZ = KFS, IGK or TVG). These constructs reproduced the expression profile specified by the GFP score in both *in vitro* and *in vivo* experiments (Fig 3D and Supplementary Fig 13). However, insertion of a 6xHistitidine tag between the second codon and the variable sequence (MV-6xHis-XYZ-eGFP), equalized expression of all constructs both *in vivo* and *in vitro* (Fig 3D and Supplementary Fig 13) arguing that the position of amino acid and nucleotide motifs drives protein synthesis efficiency.

To demonstrate that the MV-KFS-6xHis-eGFP protein had the same properties as the WT eGFP protein, we purified both proteins and analyzed their spectral properties (Supplementary Fig 14). Addition of three amino acids (KFS) and a 6xHis-tag increased overall protein production but did not change either quantum yield (Q_SKG_=0.72 for WT eGFP, and Q_KFS_=0.71 for MV-KFS-6xHis-eGFP) or absorbance spectra of eGFP variants^41^. Introduction of the same motifs in the mEOS2 coding sequence reproduced the same expression profile as previously determined expression scores for KFS, IGK and TVG motifs (Fig 3E). The nucleotide and amino acid sequence of photoconvertible fluorescent protein mEOS2 does not resemble that of eGFP, specifically in the first 5 codons (MSAIK vs MVSKG, scores of 2.79±0.41 and 2.57±0.21, for mEOS2 and eGFP respectively). Insertion of the high expressing KFS and KIH motifs in position 3-5 of the N-terminally 6xHis tagged human Gα_i_ protein (hGα_i_) resulted in significantly increased expression of recombinant protein (Fig 3F). The two proteins, mEOS2 and hGα_i_, were expressed from pBAD (Invitrogen) and pET16b vectors (Novagen) which contain different promoters (ARA vs T7 promoter, respectively), 5’ untranslated sequences (UTRs) and even different number of nucleotides between ribosome binding sites (RBS) and start codons (12 vs 7, respectively). These data argue for the significant effect of nucleotide sequence at position +7 to +15 and amino acid sequence at position 3-5 on efficiency of protein synthesis regardless of either upstream or downstream sequences. Finally, we wondered if the GFP scores could predict the expression level of recombinant human protein with 4 alternative start sites in human RGS2 protein (hRGS2)^42^. *In vitro* expression of each hRGS2 variant with a C-terminal his-tag and a single starting Met-codon (M1, M5, M16 and M33) followed the previously established distribution of GFP scores (Supplementary Fig 15). M1 variant was expressed the least (GFP score 2.93), followed by medium level expression of M5 (3.52) and M16 (3.45) variants, and highest expression of M33 variant (4.01). Together, these data demonstrate that tested motifs have a strict positional bias (nucleotides 7-15, amino acids 3-5) and are able to modulate protein synthesis efficiency regardless of the differences in the vector (promoter, terminator), upstream non-coding sequence (5’UTR and RBS) or downstream coding sequence (eGFP, mEOS2, hGαi or hRGS2).

Our *in vitro* and *in vivo* assays as well as our experimental data with different proteins and vectors (Fig 3 and Supplementary Fig 9-15) indicate striking differences in translation efficiency that is driven by nucleotide sequence at the 7-15 positions and that of the amino acid at the 3-5 positions of the open-reading frame. The position of the randomized sequence in our results (Fig 1-3, Supplementary Fig 2-15) as well as the previous studies ^7,8,11,12,14,16,21,25^ indicated that translation initiation or early elongation steps could be influencing efficacy of protein expression. To address this possibility, we assayed the efficiency of initiation complex formation and kinetics of peptidyl transfer using a well-defined *in vitro E. coli* translation system^43^. We did not observe any significant difference in the formation of translation initiation complex on 45 nucleotide long messages, resembling either WT eGFP (MVSKG, 13% initiation efficiency), one of the preferred AAVTAT motifs (MVKYQ, 15% initiation efficiency) or permuted (MVYKQ, 18% initiation efficiency). However, the yield of protein synthesis from the three initiation complexes varied significantly Fig 4A). While the full length MVSKGK peptide could hardly be observed after 5 minutes of incubation with ternary complexes and EFG, the MVKYQK peptide was readily detected only after only 10 seconds of incubation. Permuted MVYKQK full length product was also detected albeit with a yield less than that seen with the MVKYQK sequence (Fig 4A). This is likely due to the differences in GFP scores between YQK (GFP Score=3.88±0.2) and KYQ (GFP Score=4.89±0.2) arguing for both nucleotide and amino-acid composition and positional bias (nucleotides 7-15, amino acids 3-5) in determining efficiency of protein synthesis. Surprisingly, translation of MVSKGK peptide (GFP Score=2.57±0.21) seemed to be aborted or stalled after the incorporation of the 4^th^ or 5^th^ amino acid (Fig 4A, MVSK and MVSKG products). Pelleting of initiation complex and experiment following kinetics of peptidyl transfer once again demonstrated previously observed differences in protein synthesis between constructs with different nucleotide and amino acid motifs in positions 7-15 and 3-5, respectively. However, the difference in the peptide synthesis was not due to the translation initiation efficiency but rather to different translation rates in early elongation steps.

**Figure 4.**
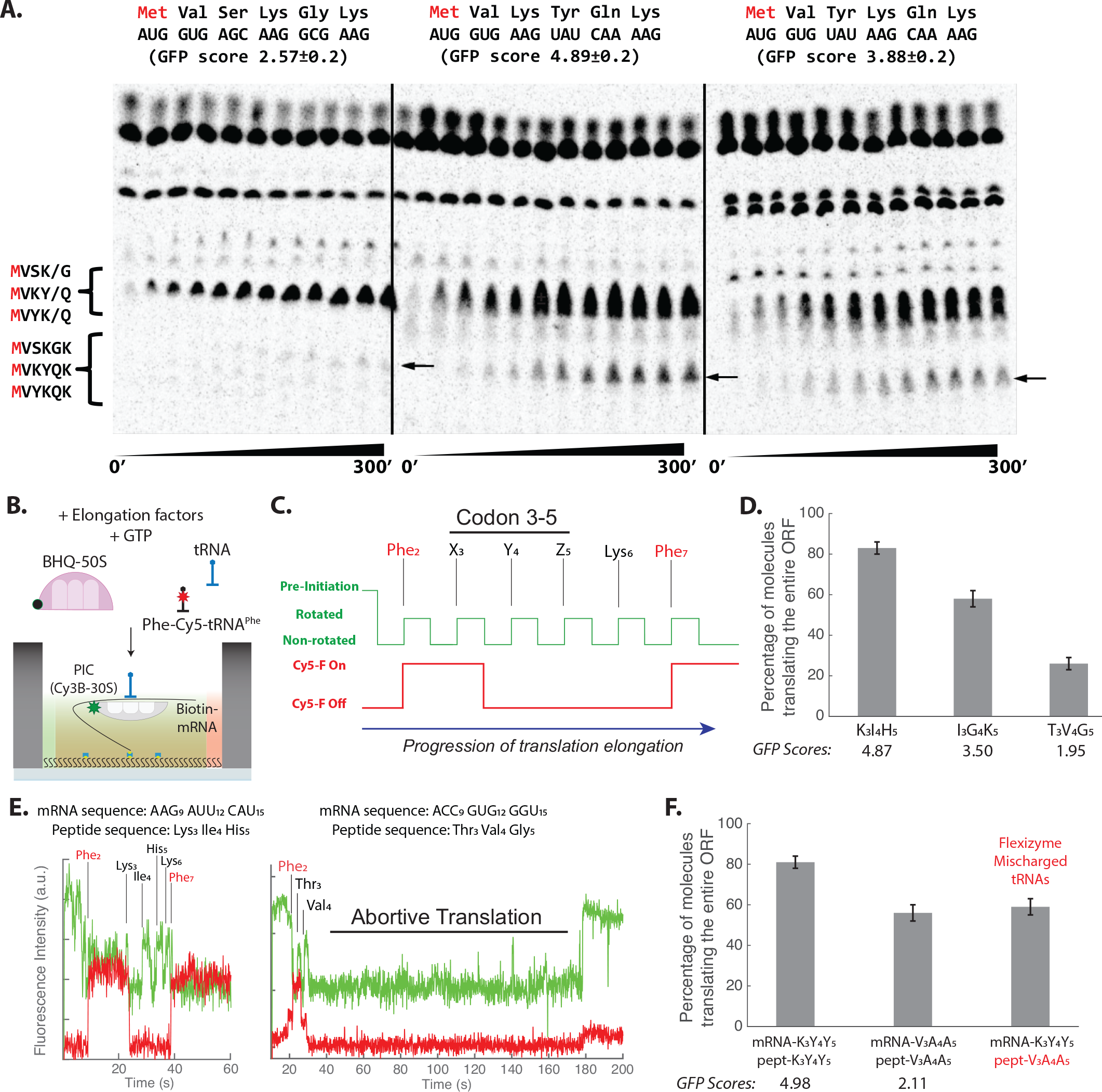
A. Thin layer chromatography (TLC) analysis of in vitro peptide synthesis using S35-labeled methionine (red). Sequences and GFP scores of penta-peptides from wild type eGFP (MVSKG) and two high expressing clones MVKYH and MVYKH are indicated. Protein synthesis was initiated at time 0 and resolved over time (300 seconds). Points at 1, 5, 10, 20, 30, 45, 60, 90, 120, 180, 240, 300 seconds are shown. Migration of tetra- and penta-peptide is indicated. Arrows indicate final hexa-peptide products of the reaction. B. Schematics of zero-mode waveguide (ZMW)-based single-molecule FRET assay to monitor translation. Elongation factors including fluorescently-labeled tRNA (Phe-Cy5-tRNAPhe) and quencher-labeled large ribosomal subunit (BHQ-50S) are delivered to pre-initiation complex (PIC) with labeled small ribosomal subunit (Cy3B-30S) and mRNA tethered to the ZMWs. C. Expected fluorescence signal observed from a translating complex utilizing FRET between Cy3B and BHQ-2 on ribosomal subunits, as well as direct excitation of Cy5 on tRNAPhe. D. Measured percentage of processively translating population for different codon 3-5 mRNA constructs (n = 179 molecules for all; error bars represent s.e. based on the binomial distribution). E. Representative traces for “processive” translation of K_3_I_4_H_5_ (Left) and “abortive” translation of T_3_V_4_G_5_ (Right). F. Measured percentage of processively translating population for K_3_Y_4_Y_5_ and V_3_A_4_A_5_ mRNA codons with a correct peptide, and for translating K_3_Y_4_Y_5_ mRNA codons with V_3_A_4_A_5_ peptide sequences using Flexizyme-mischarged tRNAs (n = 161, 147 and 156 molecules from left to right; error bars represent s.e. based on the binomial distribution).

To investigate how the resulting changes in overall translation efficiency is related to rates and processivity of translating individual codons, we used single-molecule Förster resonance energy transfer (smFRET)-based assay to monitor translation of 8-14 codons within the mRNA open reading frame (Fig 4B,C; Supplementary Figure 16) ^44,45^. Our assay monitors ribosomal conformational changes coupled to a repeated translation-elongation cycle, as well as binding of specific cognate tRNA to translating ribosome using a zero-mode waveguide-based (ZMW) experimental platform ^46^, quantifying rate and processivity for each translated codons. Ribosomal conformational changes were monitored by site-specifically labeling small and large ribosomal subunits with Cy3B fluorophore and BHQ-2 quencher, respectively. For translating each codon, a rate of tRNA-binding step during decoding and a rate of EF-G-binding step during translocation were measured as respective non-rotated and rotated ribosomal conformation state lifetimes, defined by the FRET-efficiency changes between Cy3B and BHQ-2 due to coupled ribosome conformation changes. In addition to the ribosome conformation signal, the progression of translation was independently followed by the binding and departure of fluorescently labeled Phe-specific tRNA (Phe-(Cy5)-tRNA^Phe^) to present Phe codons within mRNA. We tested three different mRNA sequences in positions 3-5 based on their expression values: K_3_I_4_H_5_ (^7^AAG AUU CAU^15^) was used for high, I_3_G_4_K_5_ (^7^AUC GGU AAG^15^) for medium and T_3_V_4_G_5_ (^7^ACC GUG GGU^15^) for low expression in otherwise identical sequence (fMet_1_-Phe_2_-X_3_-Y_4_-Z_5_-Lys_6_-Phe_7_-STOP_8_, Fig 4C). Comparing translation of high (K_3_I_4_H_5_) and low (T_3_V_4_G_5_) expression mRNA construct, we have observed a substantial alteration in translation elongation processivity, defined as the percentage of ribosomes that translated the entire ORF (Fig 4D; Supplementary Figure 17). The majority of ribosomes translating K_3_I_4_H_5_ construct completed synthesis of the peptide (Fig 4E; six cycles of elongation with two Phe-(Cy5)-tRNA^Phe^ binding events to decode F_2_ and F_7_ codons) with 84% of ribosome translating the entire ORF (Fig 4D). However, T_3_V_4_G_5_-translating ribosomes resulted in aborted protein synthesis (Fig 4E), with only 27% ribosome reached the in-frame stop codon and with a majority of translating ribosomes arrested after the incorporation of 3^rd^ (T) and 4^th^ (V) amino acid (Fig 4D). The experiment with I_3_G_4_K_5_ construct revealed intermediate ribosome processivity (54% of ribosomes translating the entire ORF, Fig 4D; Supplementary Figure 17) with bulk of translation aborted at amino acids 3 (I) and 4 (G) similar to T_3_V_4_G_5_ construct. For ribosomes that passed the “processivity barrier” at amino acids 3 and 4, both non-rotated and rotated state lifetimes for codon 3-7 were comparable across different mRNA constructs (Supplementary Figure 17), indicating possible existence of irreversible branch-points to abortive translation during the first 5 codons. The low processivity of translation observed for T_3_V_4_G_5_ construct was readily replicated in additional experiments performed at a different experimental temperature as well as at a different translational factor concentration (Supplementary Figure 18). Taken together, these data demonstrate that the abortive translation elongation via potential ribosome arrest and drop-off events at codon 3-5 is responsible for the translation efficiency differences observed in our study.

To understand a relative contribution of a peptide sequence at codon 3-5 compared to their respective nucleotide sequence in determining the overall translation efficiency, we employed specifically mischarged tRNAs to change the nascent peptide sequence without altering the mRNA sequence. Among different tripeptide sequences on codon 3-5, K_3_Y_4_Y_5_ and V_3_A_4_A_5_ were chosen due to its respective high and medium GFP score, as well as the availability of mischarged-tRNA reagents. When tested on the previously described single-molecule assay, translation of K_3_Y_4_Y_5_ was highly processive (81% of ribosomes translated the entire ORF, Fig 4F), whereas translation of V_3_A_4_A_5_ exhibited an intermediate ribosome processivity (56%, Fig 4F). tRNA^Lys^ and tRNA^Tyr^ were purified and respectively mischarged with Valine and Alanine amino-acids using flexizyme reaction ^47^, and used to translate K_3_Y_4_Y_5_ mRNA construct. Surprisingly, only changing nascent-peptide sequence to V_3_A_4_A_5_ altered the processivity of translating K_3_Y_4_Y_5_ codons to 59% (Fig 4F), similar to that of translating V_3_A_4_A_5_ codons. Our result shows that the codon 3-5 amino-acid identities contribute to the overall translation efficiency in conjunction with the mRNA sequences on position 7-15.

To gain a further insight of a structural state of ribosomes that aborted protein synthesis at codon 5, we tested whether incoming aa-tRNA can access and stably bound to the A site. As in previous experiments, we have used Cy3B signal to monitor conformation of translating ribosome and binding of Cy5-labeled Lys-tRNA (Lys-(Cy5)-tRNA^Lys^) while translating I_3_G_4_K_5_ nascent-peptide sequence (Supplementary Figure 19). Analysis of 441 smFRET traces indicated three classes of translation events (Fig 4E): Complete translation of ORF (54%), aborted translation after 4^th^ amino acid (G_4_) without Lys-(Cy5)-tRNA^Lys^ sampling (defined as tRNA binding longer than >100 millisecond ^48^ the A-site Lys codon (45%), and one that exhibited Lys-(Cy5)-tRNA^Lys^ sampling in aborted translation (1%). Considering that a majority of arrested ribosomes exhibited a non-rotated-like conformational state without (>100ms lifetime) A-site tRNA sampling necessary for a processive elongation, we hypothesize that the ribosome is in a non-canonical structural state that cannot make a stable interaction among rRNA monitoring bases and codon-anticodon duplex necessary for further elongation ^48^. Such state may be a result of different pathing of an mRNA as well as a nascent-peptide molecule within the ribosome, possibly similar to the previously observed interaction among the ErmCL nascent-peptide, the ribosome exit tunnel and the antibiotic erythromycin ^49^.

In summary, we show that the efficiency of protein synthesis in addition to overall mRNA structure and codon content is strongly dependent on the nucleotide sequence positions 7-15 and the resulting protein amino acid positions 3-5. The expression levels of 213,708 eGFP variants with randomized nucleotide and amino acid motifs in those positions resulted in four order of magnitude difference in fluorescence and protein levels. The effect of assayed motifs was dependent on both nucleotide and amino acid sequence, which suggests that a combination of tRNA, mRNA, ribosome and nascent polypeptide chain interactions define the efficiency of protein synthesis at the very N-terminus. We found that the presence of the 9 nucleotide or 3 amino acid motifs, in the correct coding position, granted better efficiency of protein synthesis for the several recombinant proteins. This was achieved regardless of their mRNA and protein sequence, the expression vector used, or *in vitro* and *in vivo* expression conditions. The different expression levels measured from the assayed sequences was not associated with efficiency of translation initiation, but instead depended on early elongation steps during translation of the 3^rd^ or 4^th^ codon. As such, the probability of the *E.coli* ribosome to synthesize N-terminal penta-peptide without ribosome arrest and abruption of translation governs the efficiency of protein synthesis. The conservation of the ribosome peptidyl-transfer center^50^, the temporal and spatial position of translation arrest and abruption, specifically during early elongation phase and enclosing first five amino acids of the nascent polypeptide chain, suggests that similar mechanism operates in other organisms as well. The motifs that we identify will assist in creating tools for higher expression of recombinant and industrial proteins as well as for further studies on how ribosomal early elongation dynamics influence protein synthesis.

## Acknowledgments

We thank members of Daniel Goldberg’s, Hani Zaher’s, David Piston’s, Joseph Puglisi’s and Sergej Djuranovic’s lab for the helpful comments. We are thankful to Philip P. Ahern, Jeffrey I. Gordon and Mikhail Berezin for assistance and equipment used in FACS sorting and quantum yield determination. This work is supported by NIH R01 GM112824 to SD, NIH R01 GM51266 to J.D.P., NIH R01 DK115972 to DWP, JDRF Award 3-APF-2018-573-A-N to ZL, Stanford Bio-X Fellowship to J.C., and NIH T32 GM: 007067 to KAC. SD holds US Provisional Patent #62/540,897 “Methods to modulate protein translation efficiency”. The authors declare that they have no competing interests.

## Methods

### Construction of Library

To create the EGFP library, the optimized EGFP sequence was amplified using primers EGFP-lib For and EGFP-Rev using Phusion – HF (NEB). The PCR product was purified using Nucleospin Gel and PCR cleanup kit (Macherey Nagel) prior to digestion with (NcoI – For) and (XhoI-Rev). The digested PCR product was ligated into digested pBAD low copy vector. The ligation product was purified using Nucleospin Gel and PCR cleanup kit (Macharey Nagel) and desalted using Illustra Microspin G-25 Columns (Thermo Fisher). The purified and desalted ligation product was then electroporated into high efficiency 5-alpha *E.coli* cells (NEB). The cells were grown overnight on LB-Carbenicillin plates and then, ~2×10^6^ colonies were scraped from the plates and collected in LB-media. An equal volume of 50% glycerol was added to the liquid culture and the cells were frozen at −80°C.

### Cell Sorting

For each FACS experiment, one vial (5 ml) of cryopreserved cells was thawed and grown in LB media with carbenicillin for 90 min. The cells were centrifuged (3000g for 5 minutes), media was removed and cells were then induced with addition of fresh media supplemented with 0.2%% L-arabinose for 3 hours. After induction, the culture was pelleted by centrifugation at 3300 g for 10 minutes and washed with PBS, followed by a second centrifugation and a final resuspension in PBS. The cells were sorted by level of GFP expression into five bins using Aria III flow cytometer (BD Biosciences) with median GFP fluorescence of 20, 120, 600, 3600 and 12,000. LB was added to the sorted cells and they were grown at 37°C for 2 hours prior to plasmid isolation using PureLink HiPure Miniprep Kit (Thermo Fisher).

### Illumina Library Preparation

PCR was performed with primers Lib_Amp_F and Lib_Amp_R and an equal mass of the plasmid isolated from each sorted bin using Phusion-HF MM (98°C for 1 min, 22 cycles: 98°C for 10 s, 55°C for 30 s, 72°C for 30 s, and 72°C for 5 min) in separate reactions. The amplicon was purified using Nucleospin Gel and PCR cleanup kit (Macherey Nagel) and then digested with NcoI and XhoI. The digested product was purified as done previously and ligated into Illumina adapters. It was then amplified using Il_Enrich_F and Il_Enrich_R using Phusion HF MM (98°C for 1 min, 21 cycles: 98°C for 10 s, 66°C for 30 s, 72°C for 30 s, and 72°C for 5 min). The product was subsequently resolved by agarose gel electrophoresis and the appropriate sized band was excised and purified. The Illumina library was multiplexed and run on four lanes of the Illumina NextSeq System.

### Sequencing analysis

Counts for each triplet codon sequence within each FACS sorted bin and the input plasmid pool were determined from our sequencing data sets. Sequences with less than 10 total counts across all bins were removed. We determined the ‘GFP Score’ by obtaining a weighted average of counts across all of the bins for a given sequence. In short, the ratio of the counts within each bin and the total across all five bins for a given sequence was multiplied by the bin number which corresponds to increased GFP expression. The average of these weighted values for each sequence was then determined to give ‘GFP Score’ :

GFP score = (Reads_bin_1/total_reads*1) + (Reads_bin_2/total_reads*2) + (Reads_bin_3/total_reads*3) + (Reads_bin_4/total_reads*4) + (Reads_bin_5/total_reads*5)

As such, 1 represents minimal and 5 maximal eGFP score; wild type eGFP sequence has a score of 2.57±0.2. We compared ‘GFP Score’ to various mRNA (GC or AT content) and peptide sequence attributes (charge and hydrophobicity) in R using custom scripts or previously described packages (peptides) respectively.

For comparison of tAI with ‘GFP Score’ we determined the tAI of all possible triplet codon sequences using *CodonR* (https://github.com/dbgoodman/ecre_cds_analysis/tree/master/codonR). To identify mRNA sequence motifs we used the R package *motifRG*. Sequences with a ‘GFP Score’ above 4 were considered ‘high’ and sequences with a ‘GFP Score’ below 3 were considered ‘low’. The same stratification was used for identifying peptides associated with mRNA sequences with high ‘GFP Score’ using the R package peplib. All scripts used for analysis and model fitting are available at the Github repository under MIT license (https://github.com/cottrellka/EGFP_library_seq).

### *In vivo* constructs expressions

Modified and wild type mEOS, eGFP and human G protein subunit alpha i1 (Giα; NM_002069) construct DNA were created by PCR using forward primers that code for the certain sequence extracted from our EGFP library expression found in the FACS experiment (KFS, KYY, KIH - high expression, IGK – moderate expression, TVG – low expression). Regulator of G protein signaling 2 (RGS2; NM_002923) constructs were amplified by PCR reaction from previously described constructs ^51^. PCR products were cloned in the pBAD or pET16b vector, transformed into Top 10 E.coli cells and sequenced for the correct clones. Correct plasmids were transformed to E*. coli* cells for *in vivo* expression (TOP10, BL21 DE3, DH5α, W3110, XAC E. coli cells were used for expression experiments). Three colonies were picked off the plates and grown overnight. Their optical density was measured and equalized to 0.1 OD at 600nm, once they reached OD_600_ of 0.5 colonies were induced with addition of L-arabinose to final 0.2% in LB-media. Expression of fluorescent proteins was followed by fluorescence normalized to number of cells. After 3 hour of induction. Same number of cells (based on OD_600_ was centrifuged and re-suspended in 2xSDS buffer. Samples were heated at 95°C for 5 minutes, after which they were frozen at −20° C for further use. Same volume of samples were loaded on 4-16% gradient SDS-PAGE gels and analyzed by western blot analysis using EGFP (JL-8; Clontech), penta-HIS (QIAGEN) or α-RF1 E.coli (Zaher Lab) antibodies. Anti-mouse or anti-rabbit HRP conjugated antibodies were used as the secondary antibodies.

### *In vitro* constructs expressions

PCR products from pBAD or pet16 cloned constructs were used as templates for NEB PURE or PUREFREX

2.*0 in vitro* translation reactions. In short, DNA constructs were amplified using Phusion – HF (NEB) kit using T7 forward primer and gene specific reverse primer. The PCR product were analyzed on agarose gels and purified using the Zymo Clean DNA gel extraction kit. Equal amounts of DNA (50-150ng) were used in *in vitro* reactions. If noted PCR products were used to synthesize RNA using T7 polymerase kit (NEB), purified using NEB RNA purification kit and equal amount of purified RNA was used for *in vitro* reaction (1 - 3ug). *In vitro* protein synthesis was conducted for 2.5 hours at 37°C if not noted differently. In case of fluorescent proteins translation was followed in parallel by fluorescence reading using for 2.5 hours in 1 minute intervals. Same amount of samples were loaded on SDS PAGE gels and western blot analyses were performed as described for *in vivo* expression experiments.

### Spectroscopy experiments

A Thermo Scientific™ Pierce™ BCA™ Protein Assay (code 10678484) has been used to have an estimate of the total protein concentration compared to a protein standard. All spectroscopic experiments have been carried out with an UV-VIS Fluorescence Spectrophotometer ISS K2. Absorbance spectrum was measured between 350 and 550 nm. Relative quantum yield is generally obtained by comparing the intensity of an unknown sample to that of a standard. The quantum yield of the unknown sample can be calculated using (ref): Q=Q_R I/I_R 〖OD〗_R/OD n^2/(n_R^2), where Q is the quantum yield, I is the integrated intensity, n is the refractive index, and OD is the optical density. “R” refers to the reference fluorophore of known quantum yield (in this case fluorescein). Since the end-point method is not accurate for the calculation of the quantum yield, we prepared solutions within the range of 0–0.01 ODs, by subsequent dilutions of the different proteins to calculate the quantum yield using the gradients determined for the sample and the reference. In this case, quantum yield is given by: Q=Q_R (Grad/ 〖 Grad〗_R)(n^2/(n_R^2)) where Grad is the gradient obtained from the plot of the integrated fluorescence intensity vs. optical density (see Supplementary Fig. 14). Absorbance and concentration of the eGFP variants was calculated for molecular weight of approximately 27kDa.

### Formation of Ribosomal Initiation Complexes

To generate initiation complexes (IC), the following components were incubated at 37°C for 30 min: 70S ribosomes (2µM), IF1, IF2, IF3, f-[35S]-Met-tRNAfmet (3µM each), mRNA (6µM) in polymix buffer containing GTP (2mM). The complexes were then purified away from free tRNAs and initiation factors over a 500µL sucrose cushion composed of 1.1M sucrose, 20mM Tris-HCl pH 7.5, 500mM NH4Cl, 0.5mM EDTA, and 10mM MgCl2. The mixture was spun at 287,000 xg at 4°C for 2 hrs, and the resulting pellet was resuspended in 1x polymix buffer and stored at −80°C. To determine the concentration of IC, the fractional radioactivity that pelleted was recorded.

### Kinetics of Peptidyl Transfer

Ternary complexes were formed as described previously ^52^. Briefly, EF-Tu (10µM final) was incubated with GTP (10mM final) and a mix of aminoacyl-tRNAs (including valine, serine, lysine, alanine, glutamine, arginine, glutamic acid, methionine, and tyrosine) in polymix buffer for 15 mins at 37°C. The ternary complex mixture was then combined with an equivalent volume of IC at 37°C. The reaction was stopped at different time points using KOH to a final concentration of 500mM. Peptide products were separated from free fMet using cellulose TLC plates that were electrophoresed in pyridine-acetate at pH 2.8 ^43^. The TLC plates were exposed to a phosphor-screen overnight, and the screens were imaged using a Personal Molecular Imager (PMI) system.

### ZMW-based single-molecule fluorescence assay to monitor translation

Overall experimental setup (using Pacific Bioscience RSII) and biological reagents have been prepared as described previously ^44–46^. Briefly, each small and large subunit were mutated to include a weakly forming RNA hairpin at helix 44 and helix 101, which was used to attach Cy3B/BHQ-2 labeled DNA oligonucleotides via RNA/DNA hybridization (labeled DNA oligonucleotides purchased from TriLink Technologies). Individual tRNA species used were purchased from Chemical Block Ltd. tRNA^Lys^ or purified from bulk E. coli tRNA ^53^. tRNA^Phe^ was labeled at acp^3^U47 position with Cy5 using NHS chemistry as previously described ^54^, with Cy5-NHS-ester purchased from GE Healthcare. Synthesis and purification of activated Ala-and Val-DBE (3,5-dinitrobenzyl esters) derivatives was done using detailed protocol ^55^. Aminoacylation of Lys-and Tyr-tRNA (Sigma Aldrich) with synthesize Val-and Ala-DBE derivatives was done using dFx ribozyme (IDT RNA oligoes) as described by Zhang and Ferre-D’Amare, 2014^56^. 5’-Biotinylated mRNAs used for single-molecule translation assay are purchased from Horizon Dharmacon. Translational factors, ribosomal S1 protein, and aminoacylated tRNAs were prepared as previously reported. All single-molecule experiments were conducted in a Tris-based polymix buffer consisting of 50 mM Tris-acetate (pH 7.5), 100 mM potassium chloride, 5 mM ammonium acetate, 0.5 mM calcium acetate, 5 mM magnesium acetate, 0.5 mM EDTA, 5 mM putrescine-HCl, and 1 mM spermidine, with additional 4 mM GTP.

Immediately before each single-molecule experiment, small and large ribosomal subunits were mixed with respective fluorescently labeled DNA oligonucleotide at 1:1.2 stoichiometric ratio in the previously described polymix buffer. Small ribosomal subunits were subsequently mixed with S1 ribosomal protein at 1:1 stoichiometric ratio, and subsequently mixed with biotinylated-mRNA, initiation factor 2, amino-acylated formyl-methionine tRNA at 1:2:13:4 in the presence of 4 mM GTP to form 30S Pre-Initiation Complex (30S PIC). 30S PIC was diluted to 10 nM in the polymix buffer supplemented with 4 mM GTP and the imaging mix (2.5 mM of PCA (protocatechuic acid), 2.5 mM of TSY, and 2X PCD (protocatechuate-3,4-dioxygenase), purchased from Pacific Bioscience; PCD added last), and incubated in the zero-mode waveguide chip treated with Neutravidin at room temperature. After immobilizing the pre-initiation complex, the chip was washed three-times using the same buffer without the complex to remove unbound complexes, and loaded onto the RSII instrument. At the same time, the delivery solution, a polymix buffer supplemented with 4 mM GTP, the imaging mix, varying concentration of tRNA ternary complexes (labeled or unlabeled), varying concentration of EF-G, and 200 nM of the BHQ-2 labeled large ribosomal subunit was prepared, and loaded onto the instrument. In general, final concentration of purified 50 nM of Phe-(Cy5)-tRNA^Phe^ (50 nM of Flexizyme-charged tRNA for applicable experiments), 0.7 µM of total delta-Phe aa-tRNA (total tRNA charged with all amino-acids except Phe; tRNA from Roche) and 100 nM of EF-G were used. Higher concentration of factors or different set of tRNAs were used as indicated in each experiments.

At the start of the experiment, the instrument delivered the delivery solution to the chip, and recorded 8-minute movie with frame rate 10 frame per second, illuminated by 60 mW per mm^2^ of 532-nm laser and 10 mW per mm^2^ of 642-nm laser. Experiments were performed with the chip temperature clamped to specified temperature, usually ranging from 20 to 30 °C. Resulting movies were analyzed using in-house-written MATLAB (MathWorks) scripts, as previously described. Briefly, traces from each zero-mode waveguide wells were manually filtered based on the presence of both fluorophores at different time points (signal from immobile fluorophores on the ribosome was expected to be present at the beginning of the movie, while signal from fluorophores attached to tRNA was expected not to be) and a single photobleaching step for each fluorophores. Filtered traces were manually assigned to rotated state and non-rotated state after the subunit joining event, cross-correlated with the labeled tRNA binding signals. From assigned traces, both rotated and non-rotated state lifetimes were calculated by fitting a single-exponential distribution to the measured state lifetimes using maximum-likelihood estimation in MATLAB.

