## Supplementary material Fig 1-19 for "Short translational ramp determines efficiency of protein synthesis"

Affiliations:

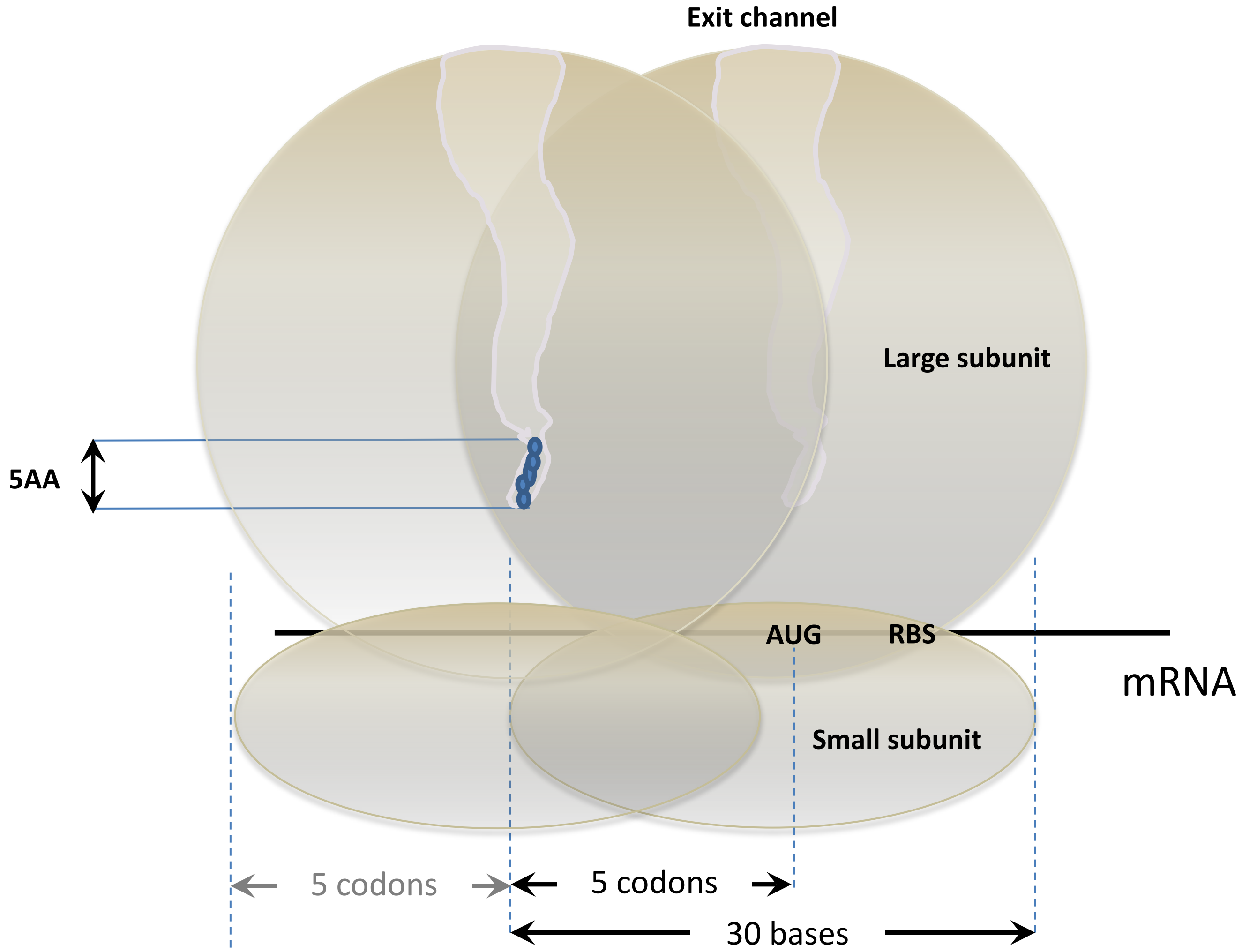


**Extended Data Figure 1 | Scheme representing first ribosome footprint and movement during the translation of the "short", five amino acid long (5AA), translational ramp described in this manuscript.** Start codon (AUG), ribosome binding site (RBS), ribosomal subunits, ribosome footprint and first five amino acids in the peptide exit channel are indicated.

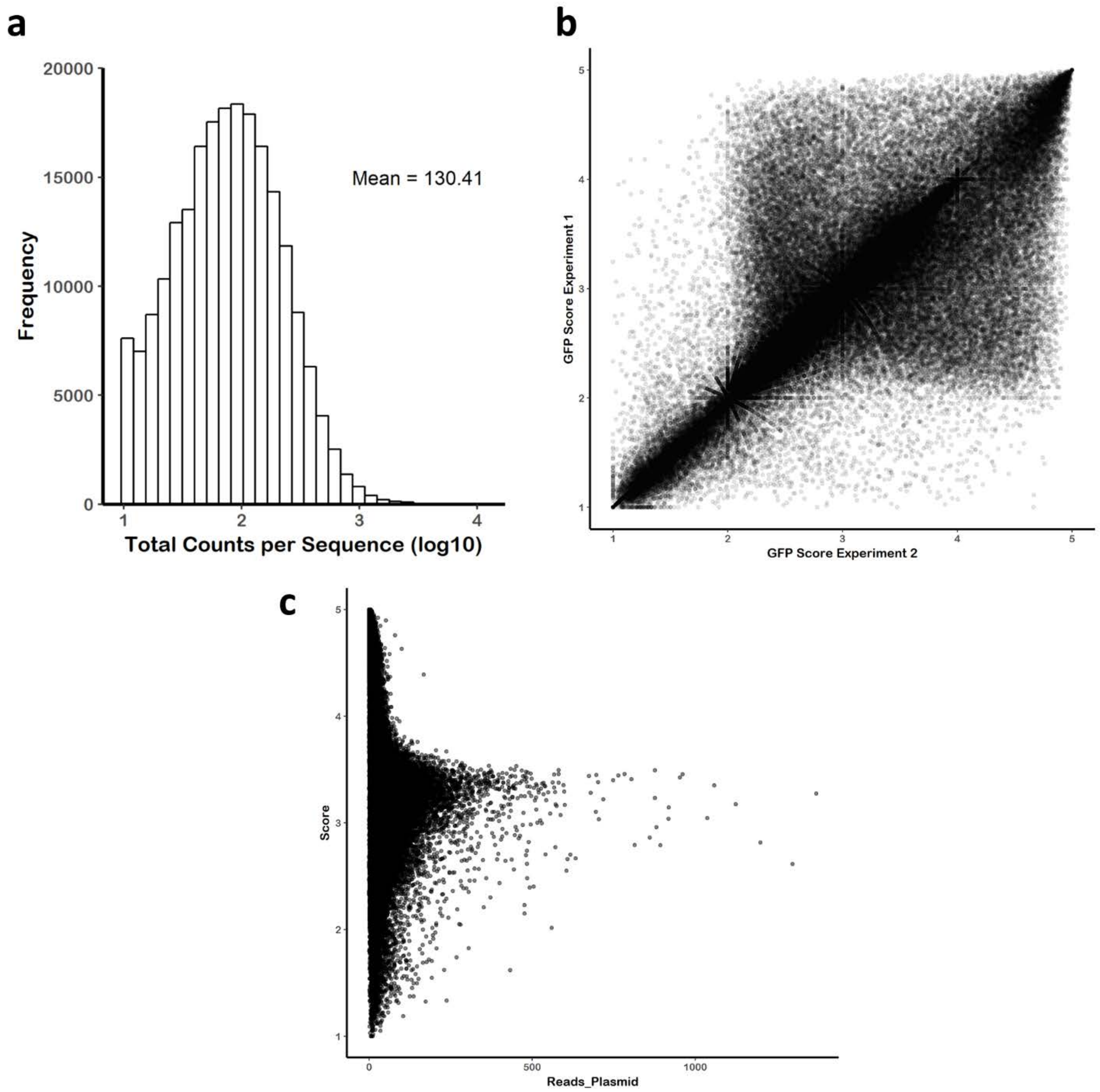


**Extended Data Figure 2 | Descriptive plots for the EGFP reporter library. a** Histogram of counts for each unique reporter construct across all five FACS sorted bins. **b** Replicate to replicate comparisons of FACS sorting experiments one and two. Pearson correlation between the two replicates is 0.796. **c** Scatter plot showing no correlation between the abundance of the reporter in the plasmid pool (Reads_Plasmid) and GFP Score.

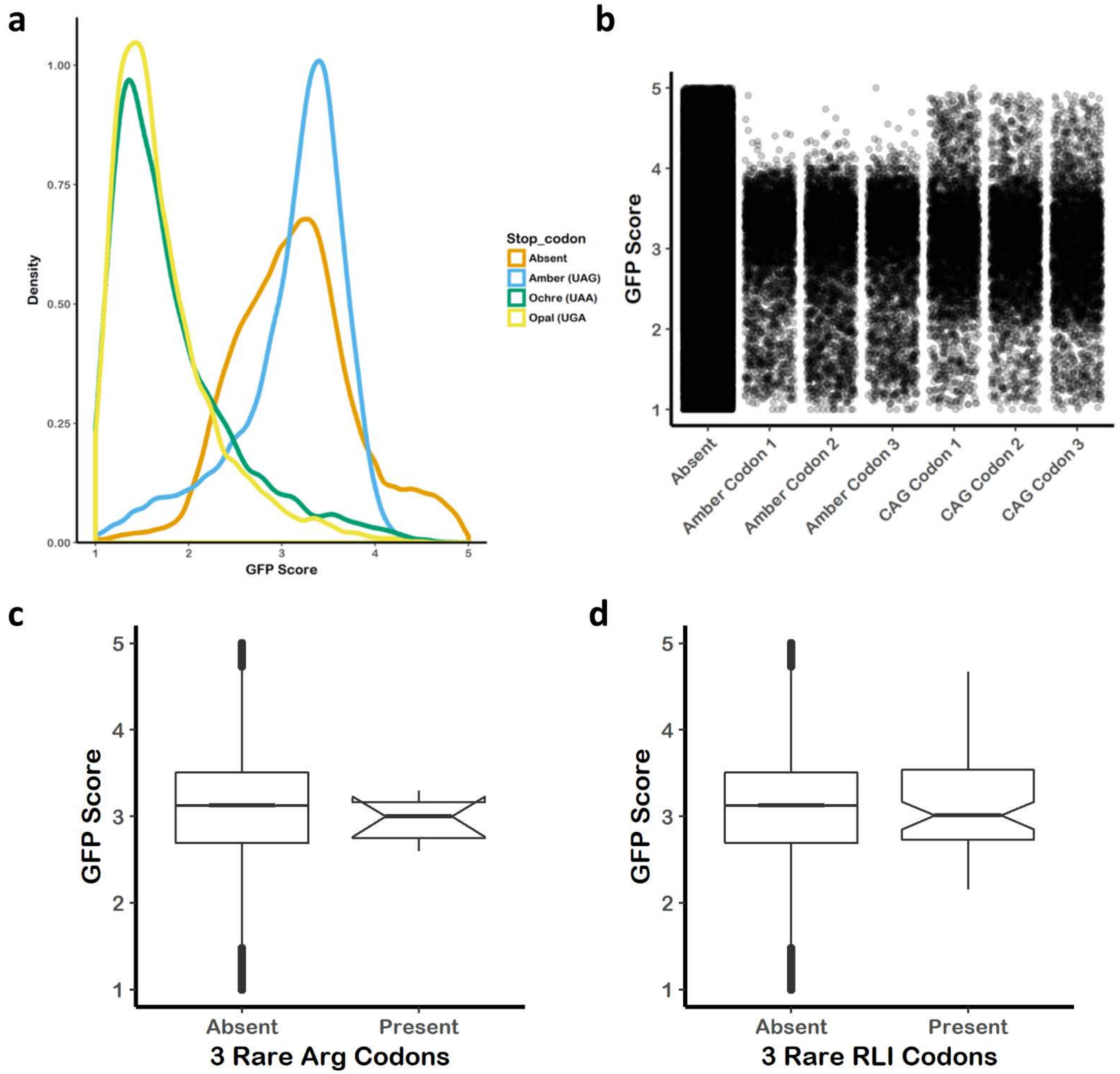


**Extended Data Figure 3 | The effect of stop codon and rare codons on GFP Score.** **a** Density plot of GFP Score for all reporter constructs lacking a stop codon (Absent) or with one or more Amber, Ochre or Opal stop codons. **b** Jitter plot of GFP Score for all constructs lacking an Amber stop codon or a $Gln^{CAG}$ codon and constructs containing either Amber or $Gln^{CAG}$ at position 1, 2 or 3 within the variable region (codons 4, 5 and 6 overall). **c** Box plot showing GFP Score for all reporters or those containing rare codons for R, L or I or R. **d** box plot showing GFP Score for reporters containing three rare arginine codons.

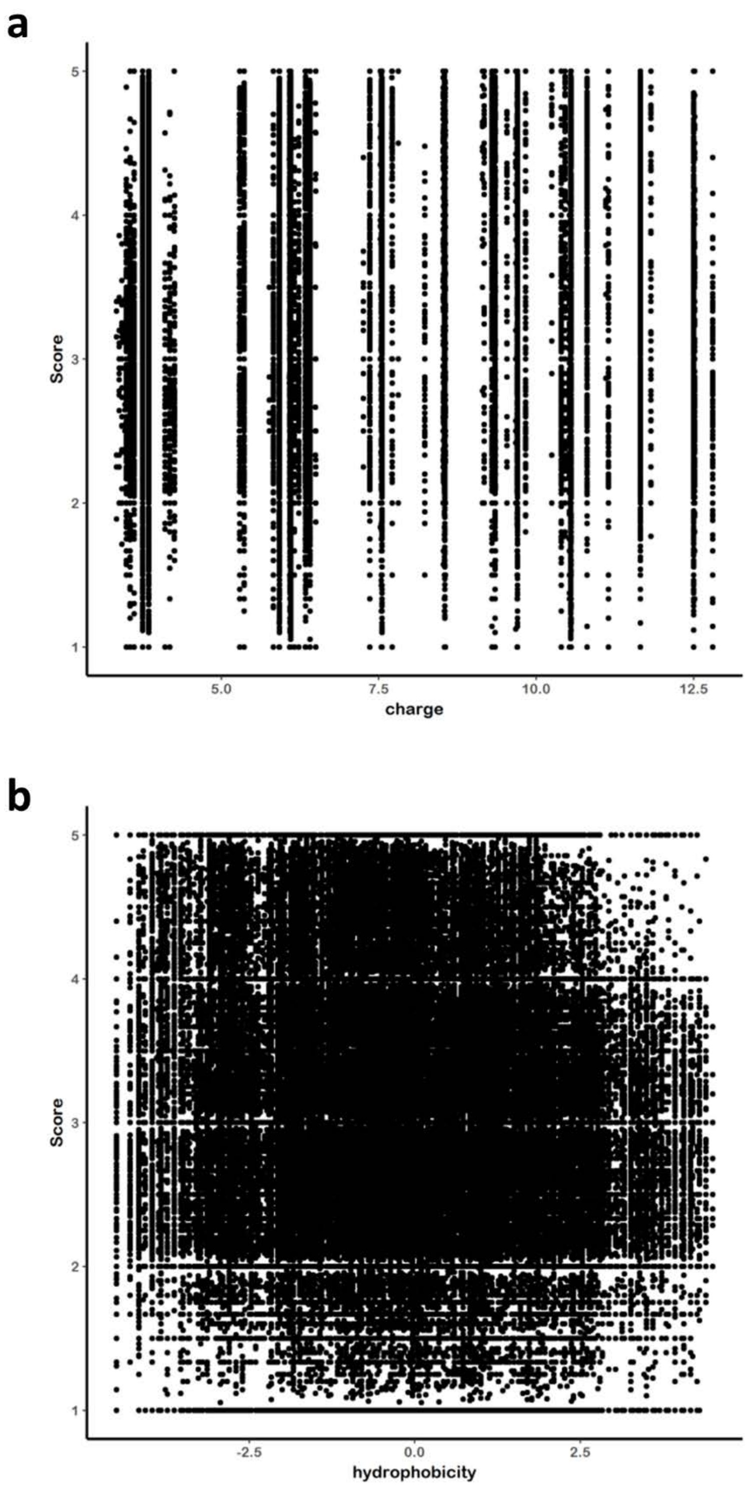


**Extended Data Figure 4 | The effective of tri-peptide characteristics on GFP Score. a** Scatter plot of GFP Score and the isoelectric point of the tri-peptide produced by the variable region within our GFP reporter library. Isoelectric point was determined by the R package Peptides using the Kyte-Doolittle scale. **b** Scatter plot of GFP Score and the hydrophobicity of the tri-peptide produced by the variable region within our GFP reporter library. Isoelectric point was determined by the R package Peptides using the EMBOSS pK scale.

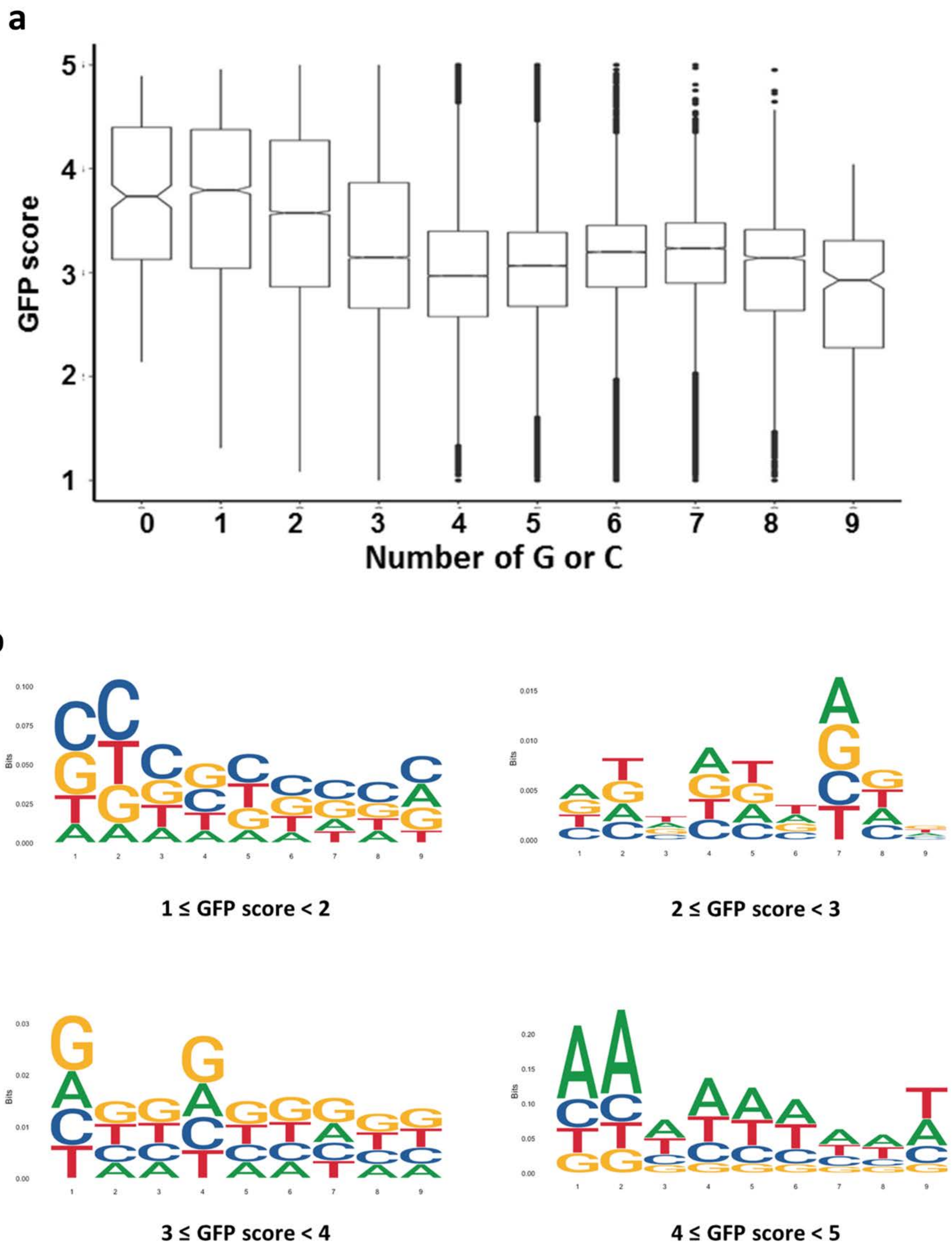


**Extended Data Figure 5 | The effect of nucleotide content on GFP Score. a** Box plot of GFP Score for all eGFP reporters within our library binned by the number of G or C nucleotides in the variable region. **B** Sequence logos for eGFP variants with defined GFP scores. Sequence logos were created from a collection of aligned sequences and depicts the consensus sequence and diversity of the sequences with each indicated score.

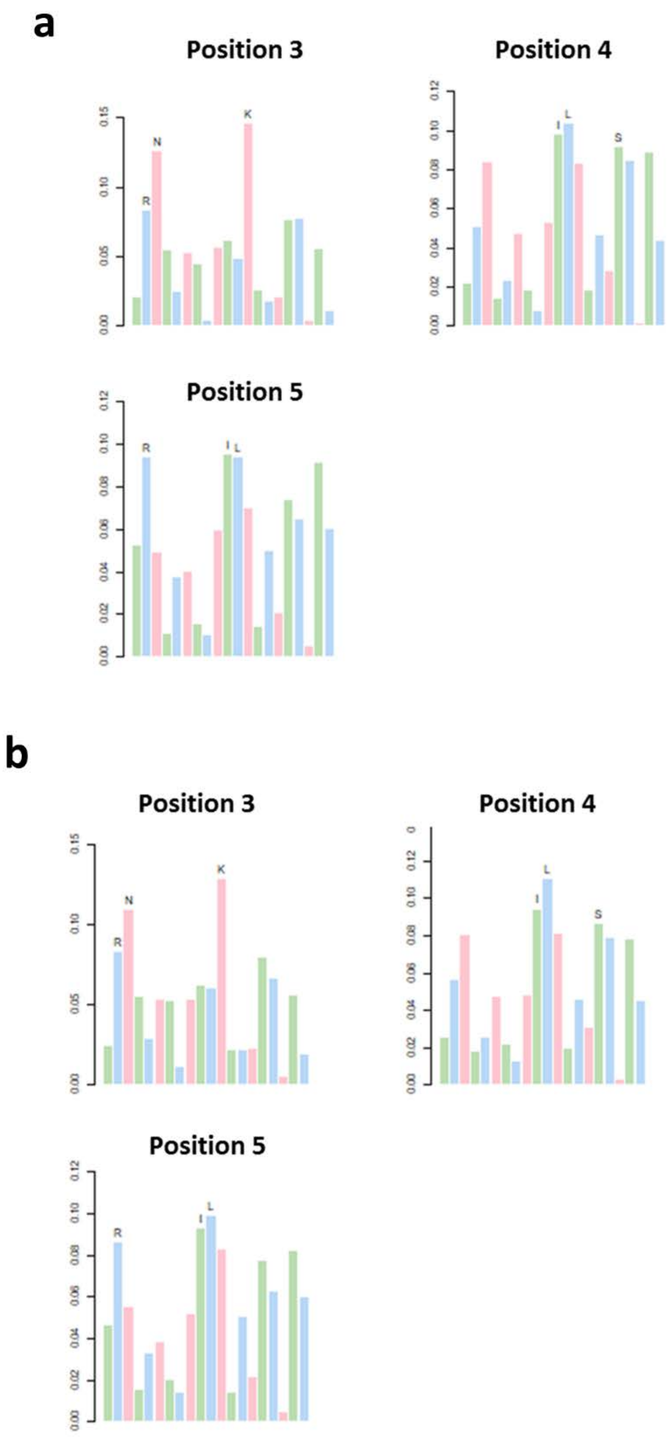


**Extended Data Figure 6 | Amino acid represntation in reporters with high GFP Score.** Bar charts showing frequency (y-axis) of each amino acid residue within the variable region of the eGFP reporters within our library for those reporters with high GFP Score (>4). Panel a shows results from experiment 1 and panel b shows results from experiment 2.

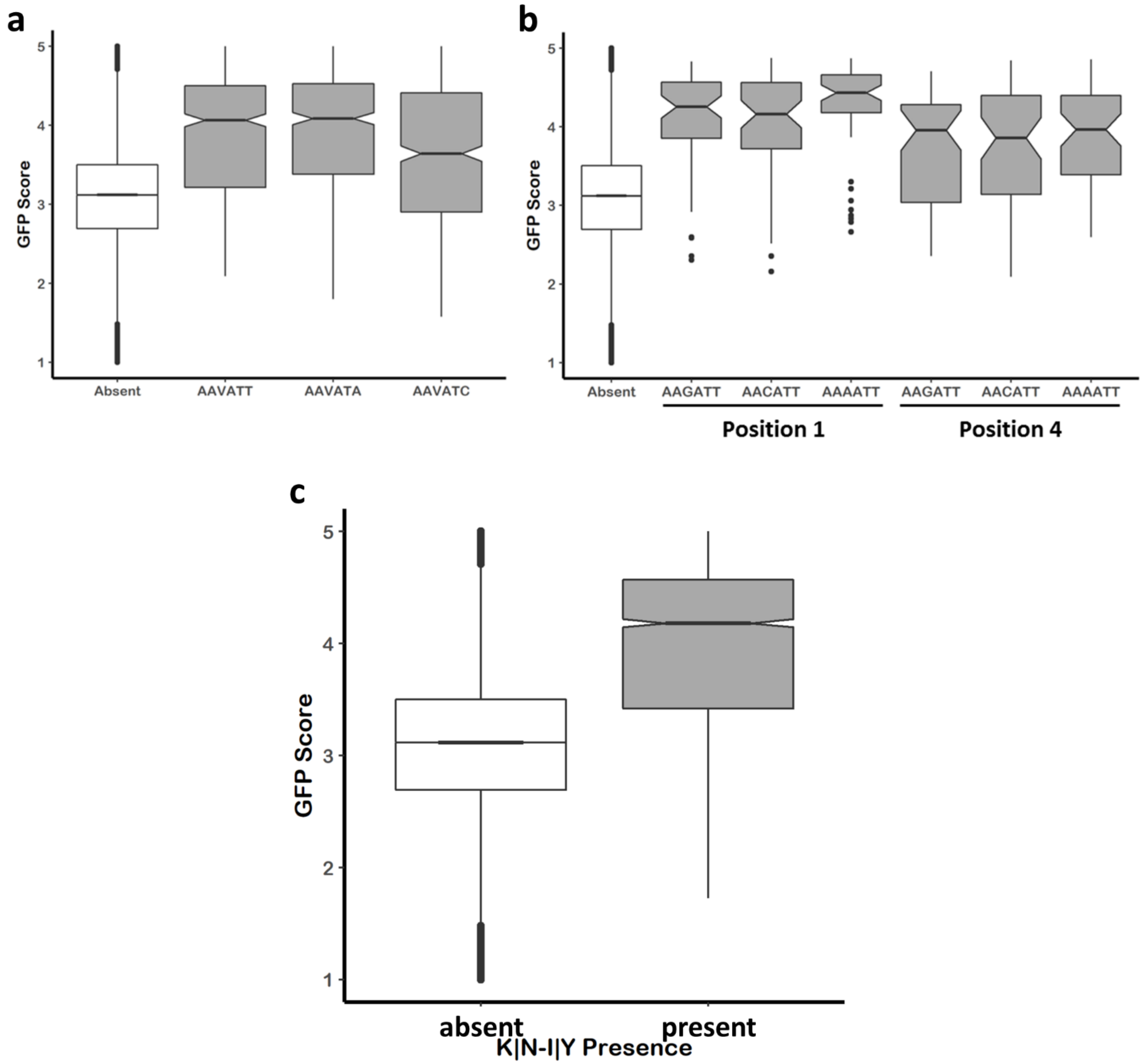


**Extended Data Figure 7 | The effect of motif variants on GFP Score.** **a** Boxplots exploring the effect of degenerate sequences of motif 1 (AAVATT) on GFP Score. **b** Boxplots comparing variants of motif 1 starting at position 1 or 4 within the variable region of the eGFP reporter. **c** Boxplots comparing eGFP reporters containing the dipeptide K or N – I or Y (KI, KY, NI, NY) within the variable region or those reporters lacking the dipeptide.

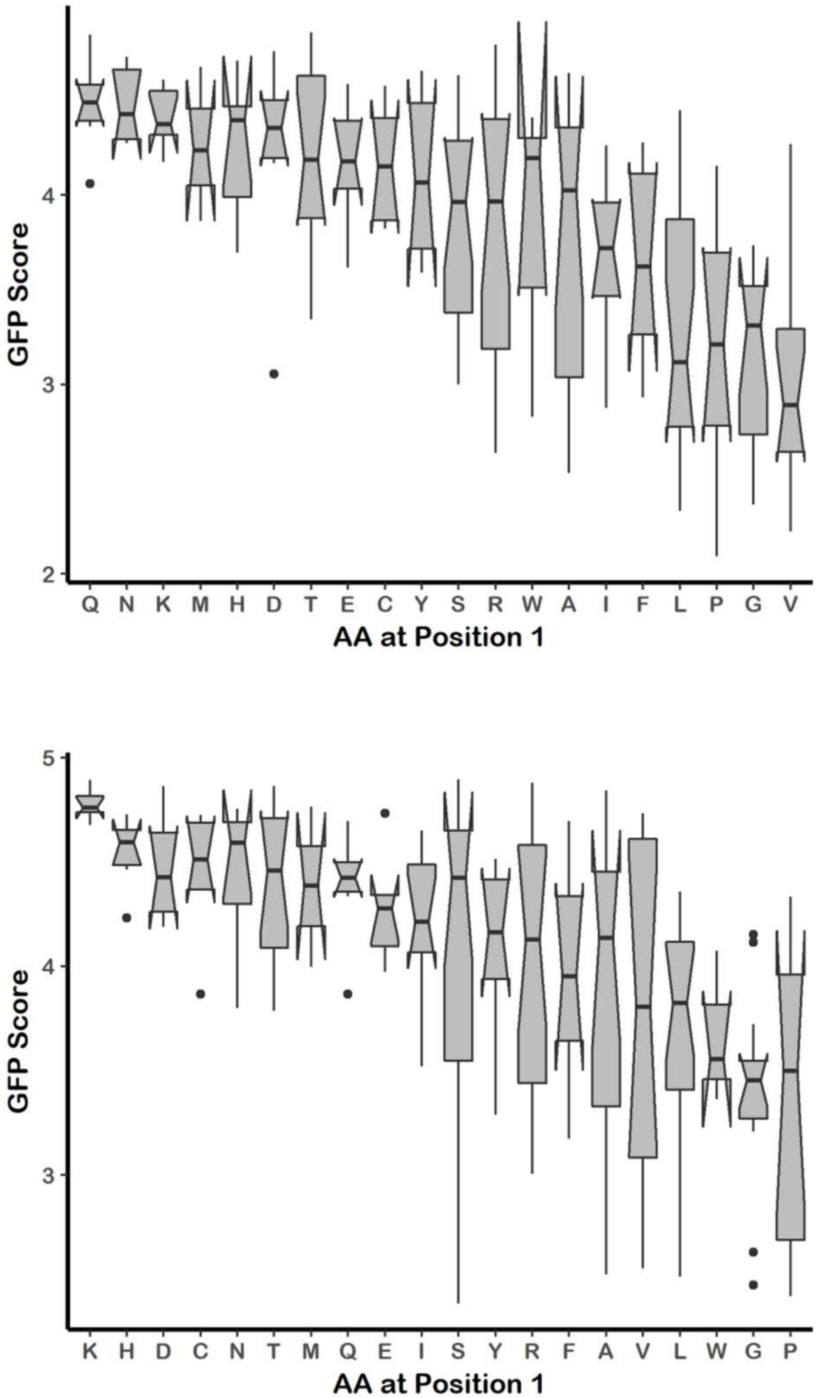


**Extended Data Figure 8 | Interplay between nucleotide motifs and the first amino acid in the variable region. a** Boxplots showing the effect of amino acid identity at position one for all reporters that contain motif 1 at position 4. **b** Boxplots showing the effect of amino acid identity at position one for all reporters that contain motif 2 at position 4.

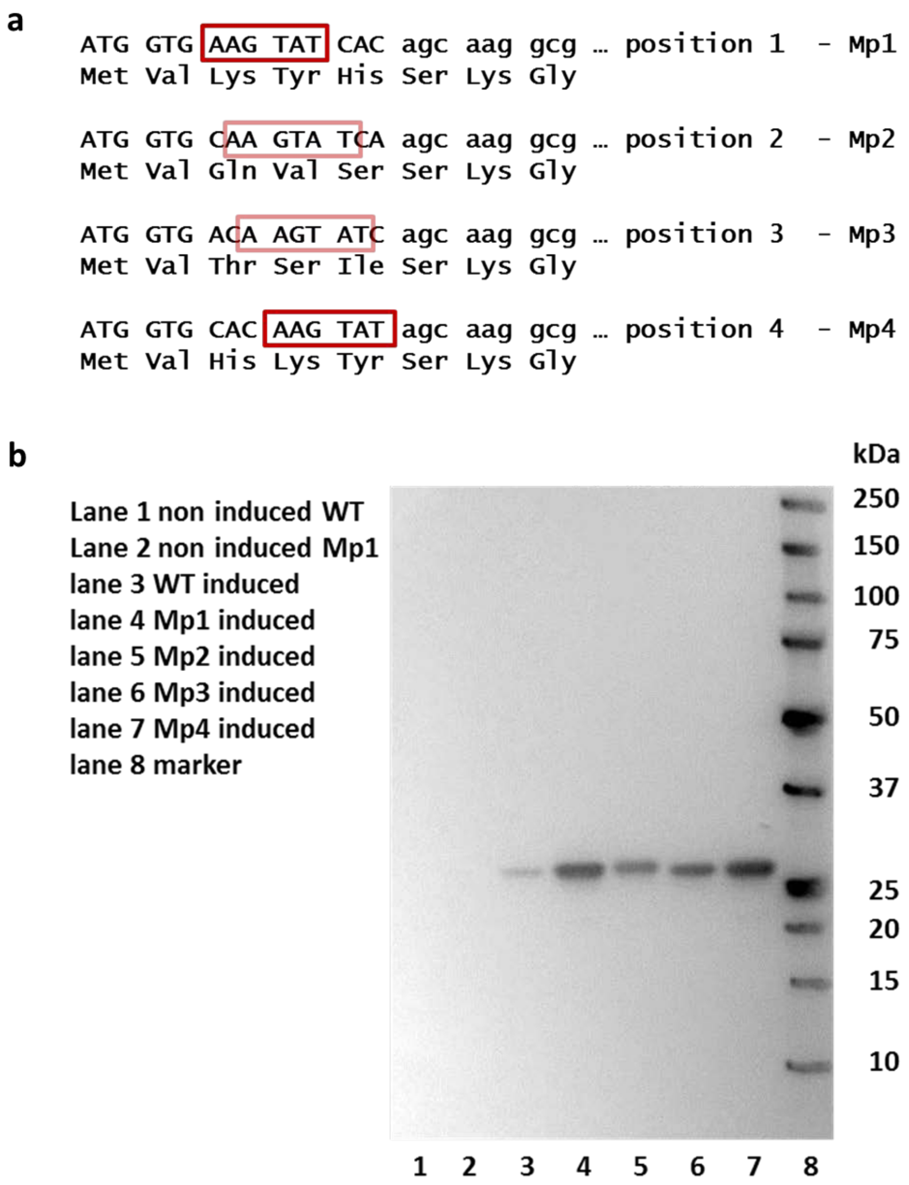


**Extended Data Figure 9 | Expression of eGFP variants with Mp1-Mp4 motifs *in vivo* confirms FACS and bioinformatics analyses. a.** Scheme of analysis of overall influence of amino acid sequence when motifs code for amino acids in positions 3 and 4 or 4 and 5, respectively. M1p1 indicates motif1 in position 1, M1p2 indicates motif1 in position 2, M1p3 indicates motif1 in position 3 and M1p4 indicates motif1 in position .Sequence of each construct and position of the motif are indicated. **b.** Western blot analysis of *in vivo* expression of eGFP constructs with motif AAG TAT in different positions coding for amino acids 3, 4 and 5. pBAD low copy vector in *E. coli* BL21 cells was used for *in vivo* expression. Equal number of E. coli cell ($OD_{600}$) following 3 hour induction with 0.2% arabinose was used for western blot analyses. Non induced wild type (WT) and Mp1 constructs are used as controls. eGFP antibody (J8, Promega) and anti-mouse HRP-conjugated secondary antibody were used to visualize expression of eGFP. Biorad Precision Plus marker is indicated in image.

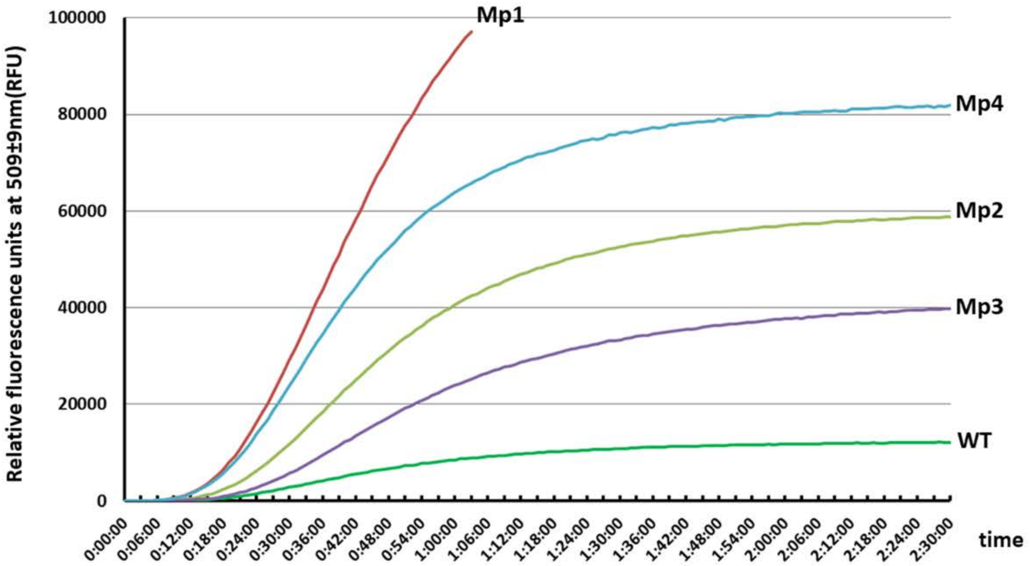


**Extended Data Figure 10 |Kinetics of of NEB Pure Express *in vitro* expression of eGFP constructs**. The expression of eGFP variants was followed by relative fluorescence (RFU) emission at 509±9nm. 100 ng of Mp1-Mp4 and WT eGFP varaints from Figure 3A-B and Extended Data Figure 9a were used as templates for in vitro reaction at 37°C.

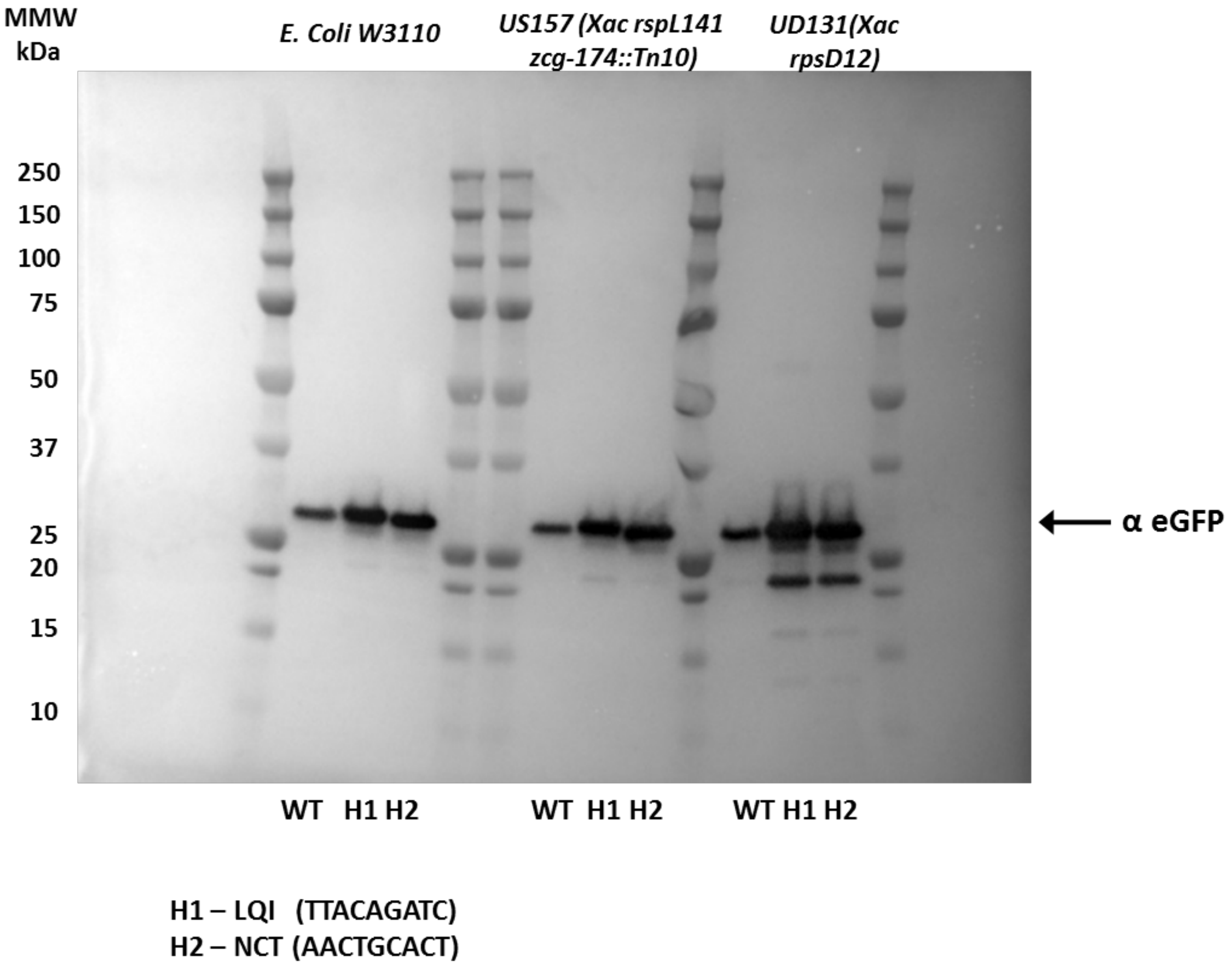


**Extended Data Figure 11 | Western blot analysis of expression of eGFP construct variants in different E. coli strains.** eGFP variants were transfected and expressed from pBAD low copy vector in *E. coli* W3110, US157 and UD131 strains. Two high expression variants H1 (NCT) and H2 (LQI) and WT eGFP constructs are indicated. Nucleotides in position 7-15 are indicated for two high expressing variants. Equal number of E. coli cell ($OD_{600}$) was used for western blot analyses after 3 hour induction period with 0.2% arabinose. eGFP antibody (J8, Promega) and anti-mouse HRP-conjugated secondary antibody were used to visualize expression of eGFP. Biorad Precision Plus marker is indicated in image.

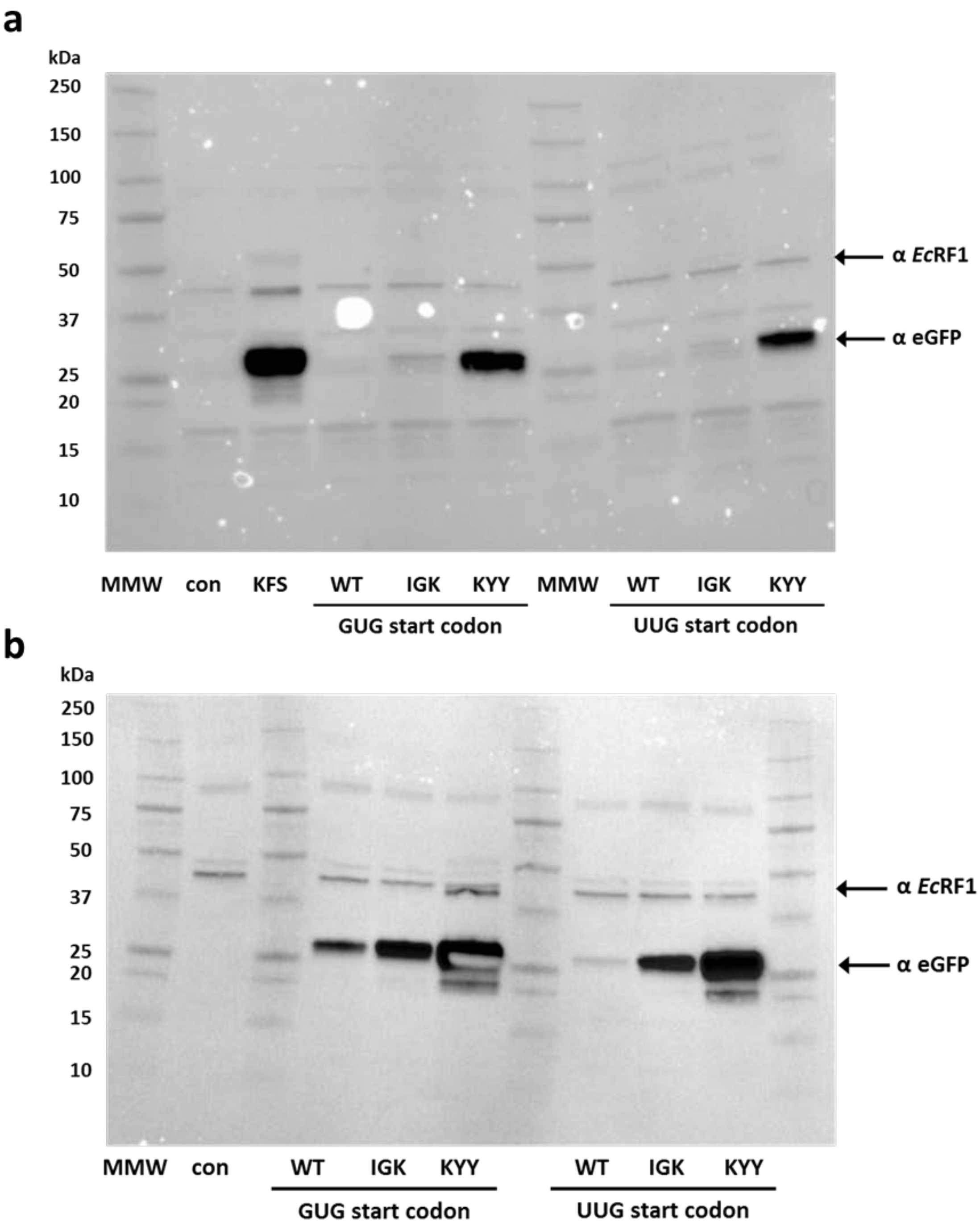


**Extended Data Figure 12 | Alternative start codons show same difference in expression of eGFP variants**. **a** Western blot analysis of NEB Pure Express *in vitro* expression of eGFP constructs with near cognate start codons GUG and UUG. eGFP construct with AUG start codon and KFS peptide in amino acid position 3-5 is indicated. 100ng of each PCR product with T7 promoter seuqnce was used for *in vitro* expression. Control (con) is NEB Pure Express *in vitro* expression reaction mix without template. 5% of in vitro translation reaction is analyzed. IGK (Gfp score 3.01±0.44) and KYY (GFP score 4.84±0.25) constructs indicate amino acids in positions 3-5 of eGFP variants. **b** Analysis of near cognate start codons GUG and UUG eGFP variants indicates previously reported difference in translation initiation efficiency (Hecht et al., 2017). Expression of GUG start codon variants is slightly increased over UUG start codon variants. eGFP antibody (J8, Promega), *E. coli* peptide release factor I (α*Ec*RF1), anti-mouse and anti-rabbit HRP-conjugated secondary antibodies were used to visualize expression of eGFP and normalization of western blot data, respectively. Biorad Precision Plus marker is indicated in image.

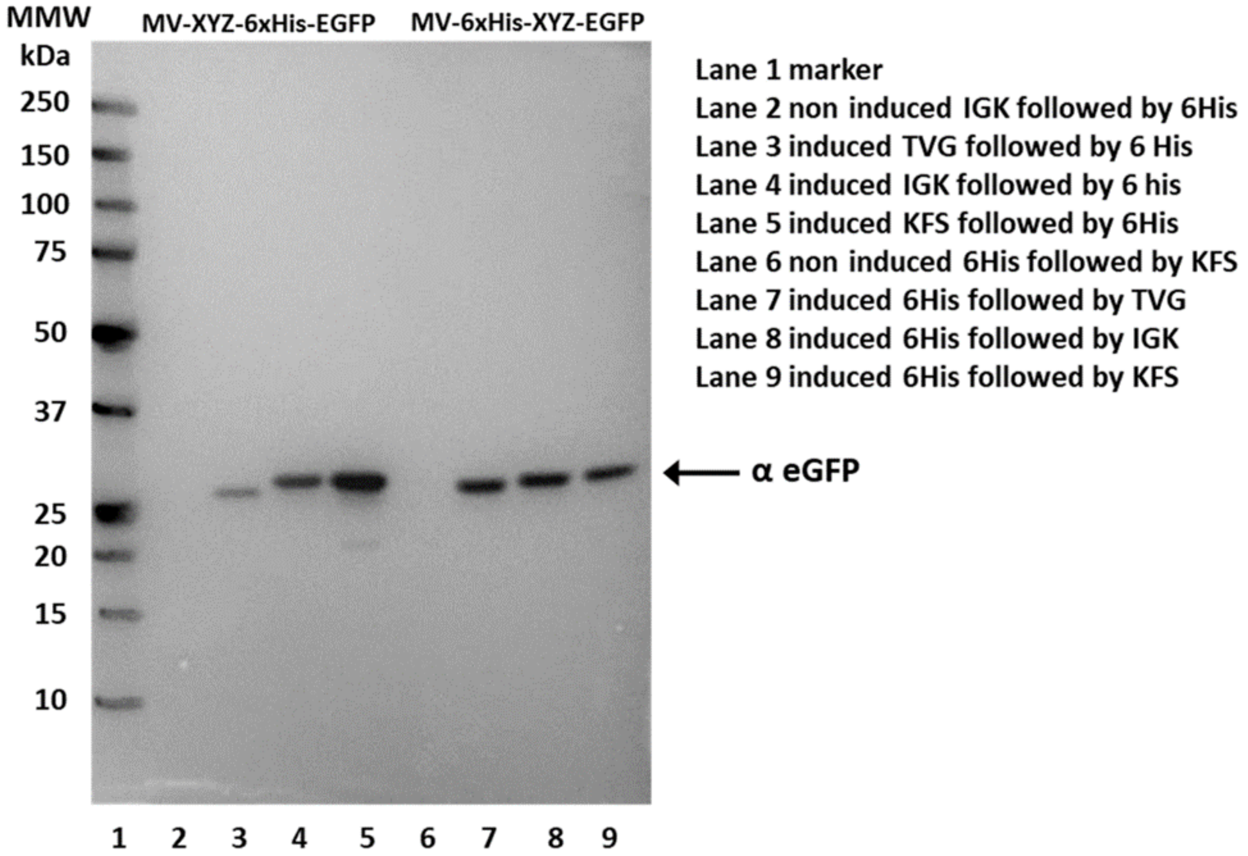


**Extended Data Figure 13 | Positional bias in controlling expression of eGFP constructs with different amino acids in position 3, 4 and 5.** Western blot analysis of *in vivo* expression of eGFP constructs with sequence XYZ (KFS, IGK and TVG, respectively) as amino acids 3(X), 4(Y) and 5(Z) followed by 6xHis tag (MV-XYZ-6xHis) or as amino acids 9(X), 10(Y) and 11(Z) preceded by 6xHis tag (MV-6xHis-XYZ). *E. coli* BL21 cells were used for expression and equal number of cells ($OD_{600}$) was used for analyses upon 3 hour induction period with 0.2% arabinose. eGFP antibody (J8, Promega) and anti-mouse HRP-conjugated secondary antibody were used to visualize expression of eGFP. Biorad Precision Plus marker is indicated in image.

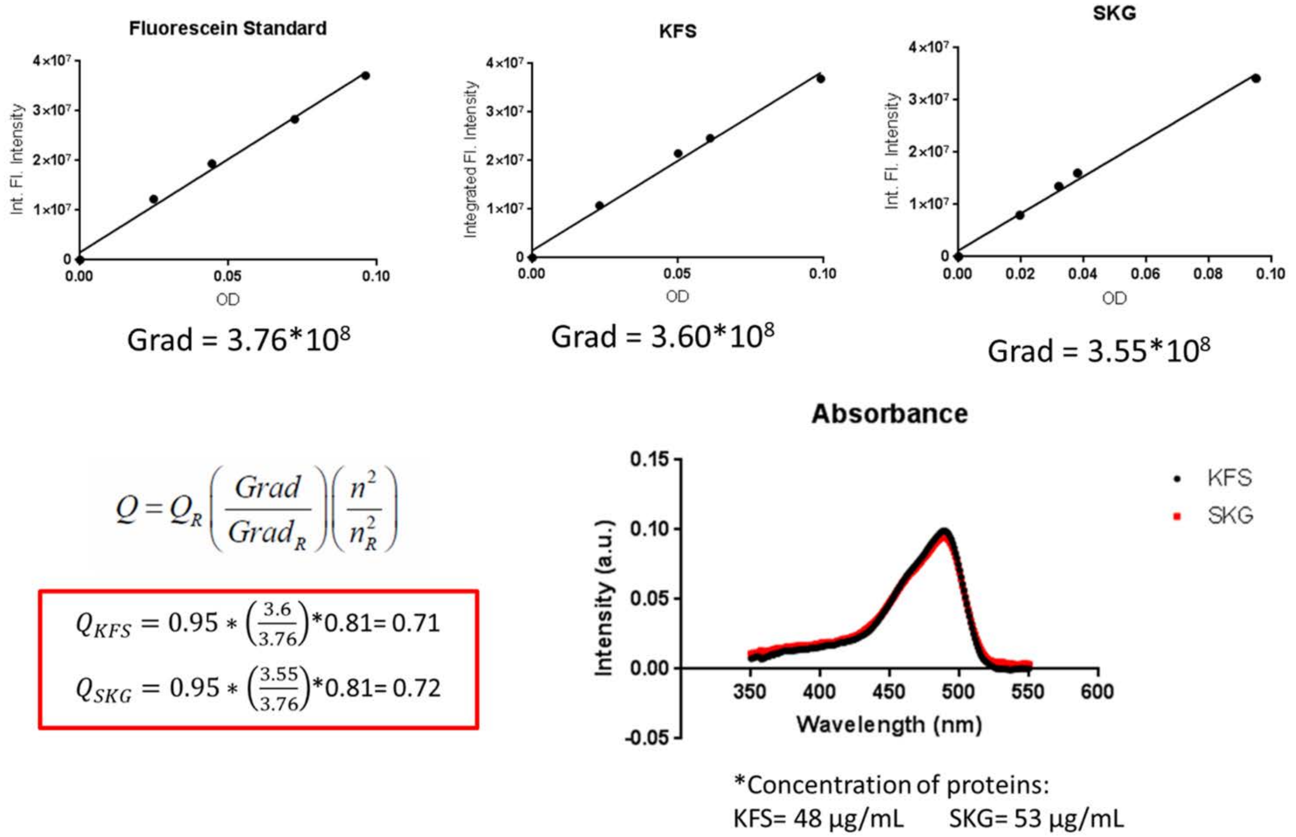


**Extended Data Figure 14 | Normalized absorbance spectrum and quantum yield calculation for the KFS eGFP variant and wt SKG eGFP.** Absorption spectra were background corrected for solvent (PBS) absorbance (lower right), and the optical density was determined as a function of excitation wavelength, in a range of values from 0.01 to 0.1. Quantum yield has been calculated as relative quantum yield compared to a fluorescein standard (Q=0.95, measured at 22°C in NaOH). The area under the curve of background-subtracted fluorescent emission spectra were calculated (Vinci ISS software) to obtain integrated fluorescence intensities (top row). The gradient (slope) values of integrated fluorescence intensity vs. optical density were linearly fit and used for the calculation of the quantum yields (red box).

```
hRGS2   MQSAMFLAVQHDCRPMDKSAGSGHKSEEKREKMKRTLLKDWKTR...
M1      MVQSALFLAVQHDCRPLDKSAGSGHKSEEKREKLKRTLLKDWKT...        (AGTGCTATG)
M5      MVLAVQHDCRPLDKSAGSGHKSEEKREKLKRTLLKDWKTRLSYR...        (TTGGCTGTT)
M16     MVKSAGSGHKSEEKREKLKRTLLKDWKTRLSYRLSYFLQNSSTR...        (AAGAGCGCA)
M33     MVKRTLLKDWKTRLSYRLSYFLQNSSTRLSYFLQNSSTPGKPKT...        (CGGACTCTT)
```

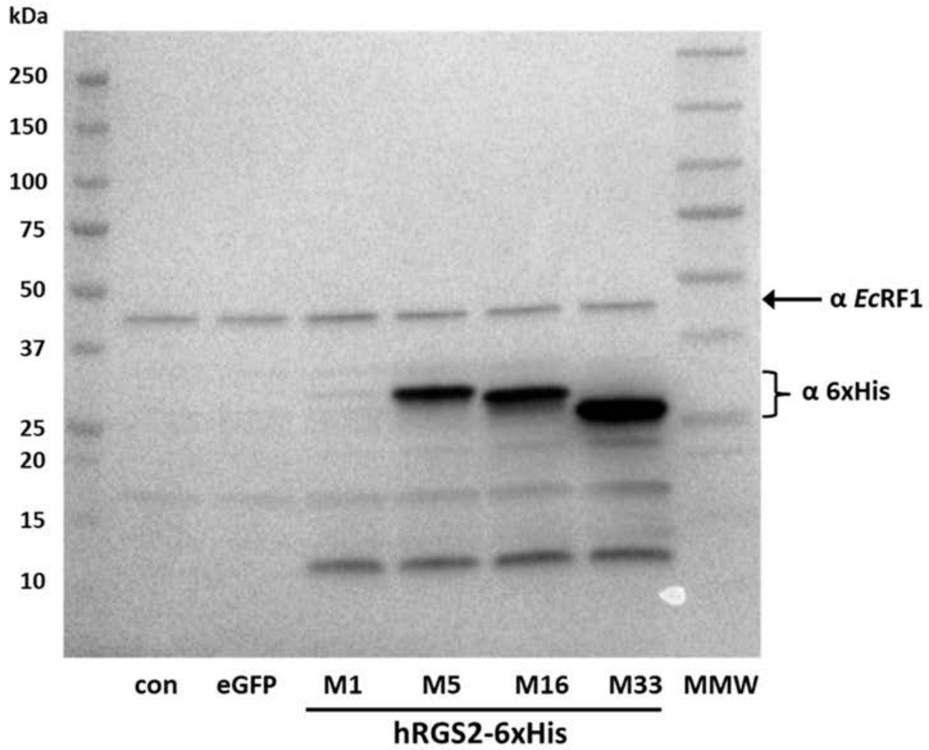


**Extended Data Figure 15 | Expression of recombinant human protein with alternative start sites in NEB PURE *in vitro* expression system follows previously obtained GFP score for eGFP varaints.** Scheme and sequences of wild type human RGS2 protein (hRGS2) and four tested constructs with alternative starting methionine (M1, M5, M16 and M33) are indicated. Alternative start site methionine (M) residues are mutated into leucines (L) to maintain one methionine residue per construct. Western blot analyses of NEB Pure Express *in vitro* expression of hRGS2 variants as well controls (con and eGFP) are indicated. 100ng of each hRGS2 variant with T7 promoter sequence was used in *in vitro* reaction. Proteins were visualized based on their C-terminal 6-Hist tag using Penta-His (Qiagen) antibody. E. coli peptide release factor I (αEcRF1) and anti-rabbit HRP-conjugated secondary antibody was used for normalization of western blot analyses. Biorad Precision Plus marker is indicated in image.

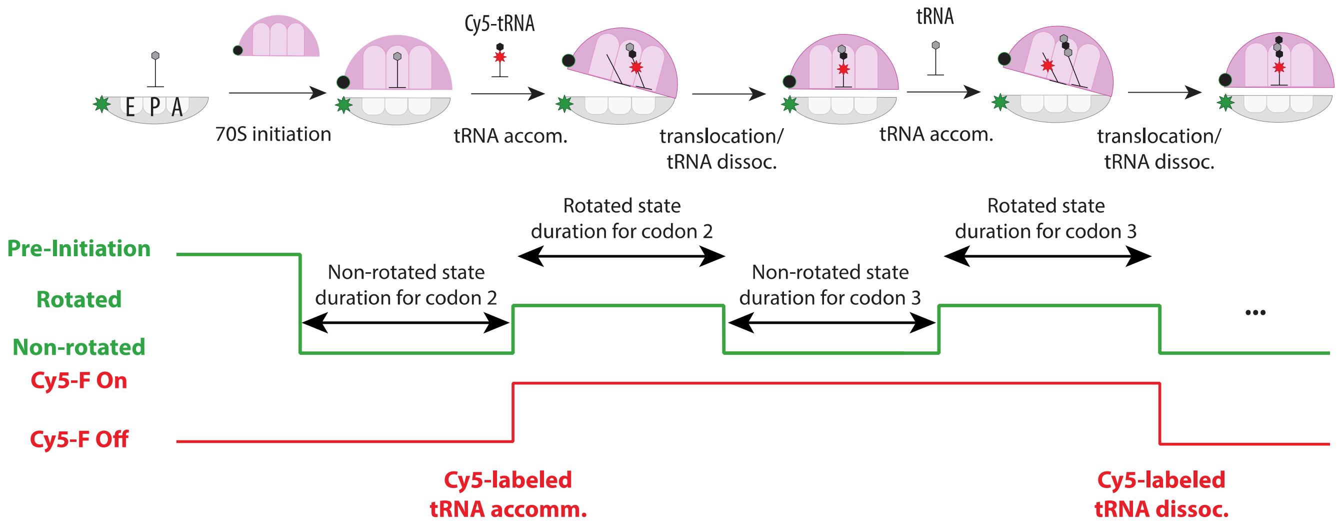


**Extended Data Figure 16 | Progression of translation elongation and expected single-molecule fluorescence signal in each step.** Prior to the experiment, Pre-Initiation Complex (PIC) composed of Cy3B-labeled small ribosomal subunit (Cy3B-30S; Cy3B denoted as a green star in the schematics), initiation factor 2 (IF2; not shown), initiator fMet tRNA and 5'-biotinylated mRNA (not shown) is formed and tethered to the SMRT cell (purchased from Pacific Biosciences) patterned with zero-mode waveguides. The SMRT cell is Neutravidin-coated to allow tethering of PIC via Biotin-Neutravidin interaction from the 5'-end of mRNA. In the beginning of the experiment, buffer containing elongation factors (including Phe-Cy5-tRNA$^{Phe}$; Cy5 denoted as red star in the schematics) and BHQ-2-labeled large ribosomal subunit (BHQ-50S; BHQ denoted as black circle in the schematics) are delivered to the SMRT cell. Upon initiation of translation, BHQ-50S binds to the 30S PIC and brings BHQ-2 close to Cy3B such that Cy3B fluorescence (denoted as green trace) is quenched by Förster Resonance Energy Transfer (FRET) at the non-rotated state. After 70S initiation step, cognate tRNA to codon presented in the A site (Phe codon in this case) can bind to the ribosome and accommodated for peptidyl-transferase reaction. Successful peptide-transfer is coupled with ribosome conformational change from non-rotated state to rotated state, where ribosomal subunits rotate respect to each other by about 7 degrees. Rotated state is a correct substrate for translocation catalyzed by elongation factor-G (EF-G; not shown), which moves the complex to the next codon and resets ribosomal conformation to non-rotated state, allowing next cycle of elongation to begin. Time between transitions are collected and fitted via single-exponential function to calculate non-rotated and rotated state lifetimes for codon present in the A site.

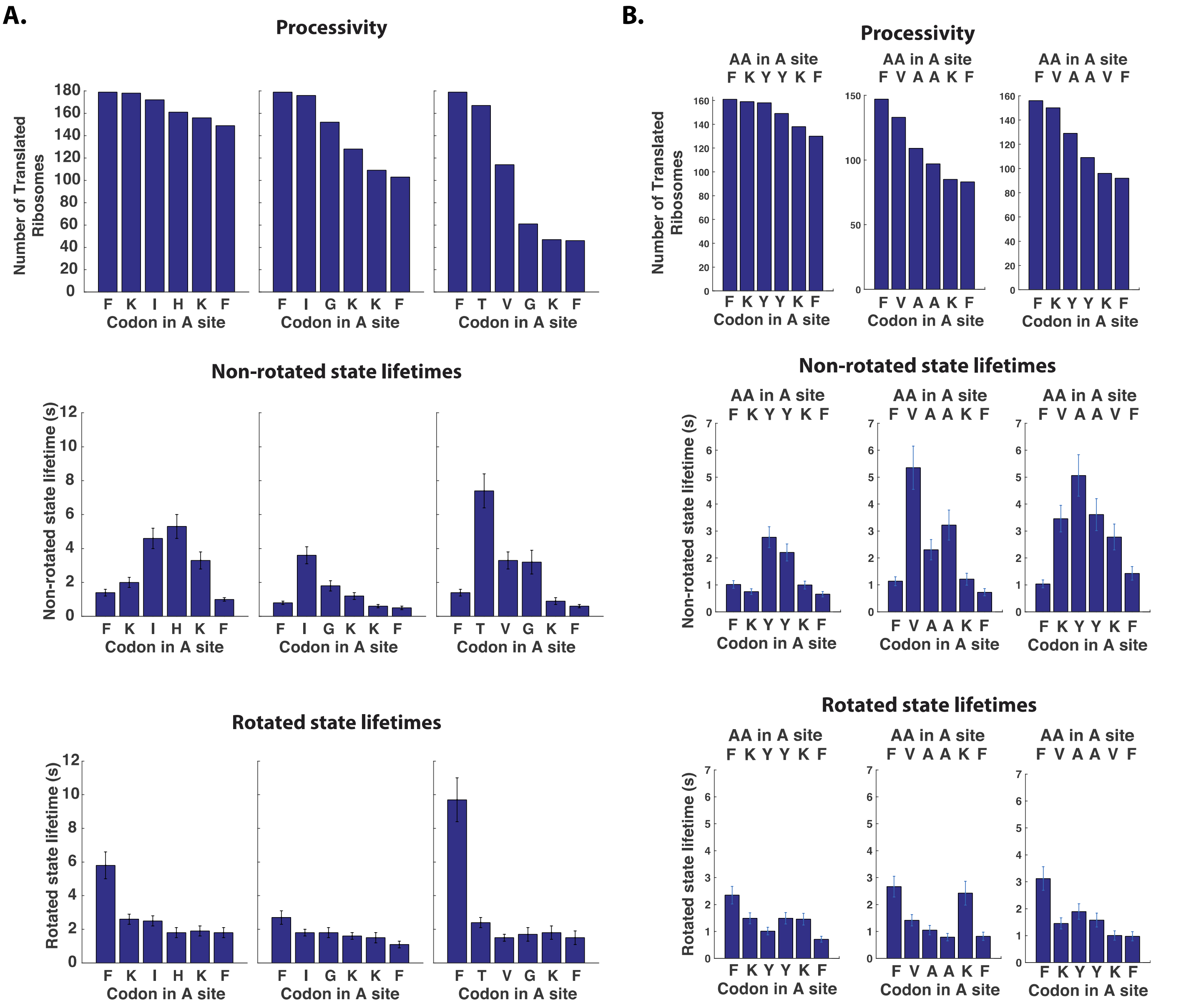


**Extended Data Figure 17 | Processivity and state lifetime measured for data shown in Figure 4D and 4F.** For each mRNA construct with different codon 3-5, the ribosome processivity as well as non-rotated and rotated state lifetimes were measured for each codons (as described in the **Extended Data Figure 16**). **A.** Processivity and state lifetime measured for construct shown in **Figure 4D**. **B.** Processivity and state lifetime measured for construct shown in **Figure 4F**. Percentage of molecules translating the entire ORF is calculated as percentage of ribosomes that have translated the last codon within the ORF ($F_7$ in this case) of the ribosomes that have translated the first codon after the start codon ($F_2$ in this case). Error bar shown in this figure is s.e. from fitting the single-exponential distribution.

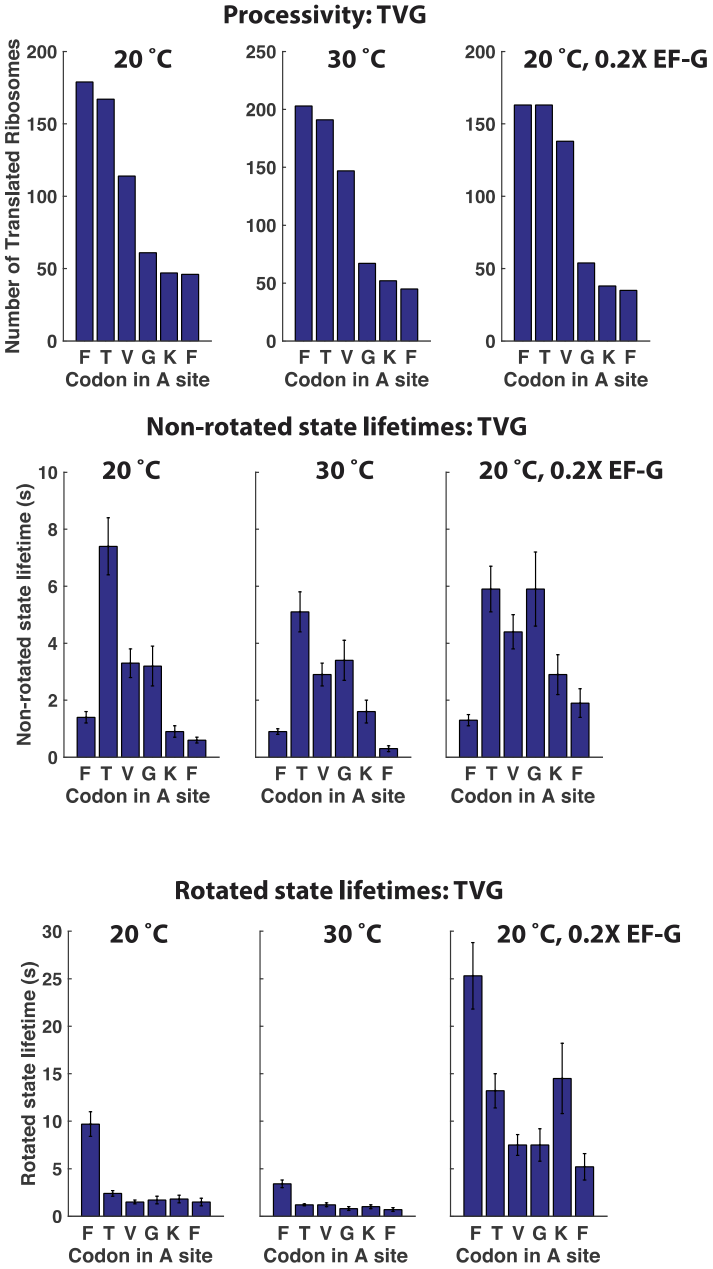


**Extended Data Figure 18 | Processivity and state lifetime measured for $T_3V_4G_5$ construct at different experimental condition.** For $T_3V_4G_5$ mRNA construct with different codon 3-5, the ribosome processivity as well as non-rotated and rotated state lifetimes were measured for each codons (as described in the **Extended Data Figure 16**) at three different conditions: at 20 °C (same as shown in the **Extended Data Figure 17**), at 30 °C, and at 20 °C with 5-fold less concentration of elongation factor-G (EF-G). The processivity shown here at top is used to calculated the percentage of molecules translating the entire ORF: 26 ± 3 % (at 20 °C), 22 ± 3% (at 30 °C) and 21 ± 3% (at 20 °C with 5-fold less EF-G; 20 nM). Lowering EF-G concentration increases the rotated state lifetimes for all codons, as they are rate-limited by EF-G association kinetics. Error bar shown in this figure is s.e. from fitting the single-exponential distribution.

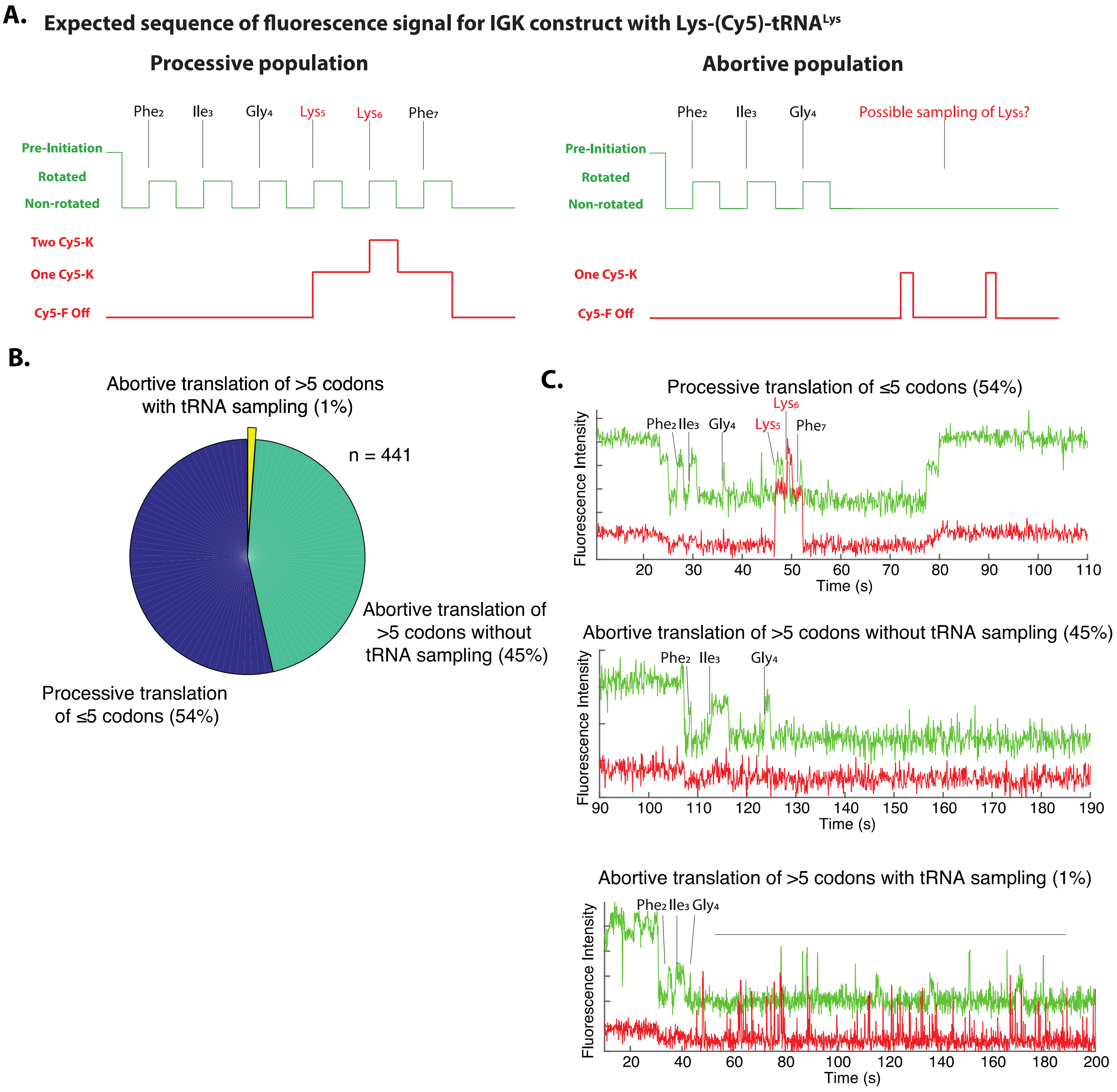


**Extended Data Figure 19 | Monitoring sampling of a cognate tRNA to the abortive translation complex at codon 5.** Using Cy5-labeled Lysine-specific tRNA (Lys-Cy5-$tRNA^{Lys}$), possible tRNA sampling of the ribosomal A site is monitored after translational arrest at codon 5. **A.** Expected fluorescence traces for successful and abortive translation, and possible sampling of tRNA to the A site on the arrested ribosmal population. **B.** A pie-chart of different populations observed in the experiment. **C.** Representative traces for each population.
